# Stress-driven transposable element de-repression dynamics in a fungal pathogen

**DOI:** 10.1101/633693

**Authors:** Simone Fouché, Thomas Badet, Ursula Oggenfuss, Clémence Plissonneau, Carolina Sardinha Francisco, Daniel Croll

## Abstract

Transposable elements (TEs) are drivers of genome evolution and affect the expression landscape of the host genome. Stress is a major factor inducing TE activity, however the regulatory mechanisms underlying de-repression are poorly understood. Key unresolved questions are whether different types of stress differentially induce TE activity and whether different TEs respond differently to the same stress. Plant pathogens are excellent models to dissect the impact of stress on TEs, because lifestyle transitions on and off the host impose exposure to a variety of stress conditions. We analyzed the TE expression landscape of four well-characterized strains of the major wheat pathogen *Zymoseptoria tritici*. We experimentally exposed strains to nutrient starvation and host infection stress. Contrary to expectations, we show that the two distinct conditions induce the expression of different sets of TEs. In particular, the most highly expressed TEs, including MITE and LTR-*Gypsy* elements, show highly distinct de-repression across stress conditions. Both the genomic context of TEs and the genetic background stress (*i.e.* different strains harboring the same TEs) were major predictors of de-repression dynamics under stress. Genomic defenses inducing point mutations in repetitive regions were largely ineffective to prevent TE de-repression. Consistent with TE de-repression being governed by epigenetic effects, we found that gene expression profiles under stress varied significantly depending on the proximity to the closest TEs. The unexpected complexity in TE responsiveness to stress across genetic backgrounds and genomic locations shows that species harbor substantial genetic variation to control TEs.

## Introduction

Transposable elements (TEs) are mobile genetic elements that were first discovered in maize (McClintock 1950) and propagate in genomes without apparent benefit to the host (Doolittle and Sapienza 1980). Uncontrolled spread of TEs is thought to have a fitness cost to the host due to the increased genome size and higher likelihood of deleterious, non-homologous recombination events (Chuong et al. 2017; Mita & Boeke 2016). TEs are subdivided into two major categories according to their mechanism of replication, namely, class I TEs that transpose through an RNA intermediate (*i.e.* RNA transposons) and class II TEs that transpose through a cut-and-paste mechanism (*i.e.* DNA transposons). Both classes of TEs are expressed. Host genomes have co-evolved with their TEs to suppress their expression (Slotkin and Martienssen 2007). These mechanisms include epigenetic silencing through histone modifications or DNA methylation, targeted mutagenesis and small RNA interference. In order to autonomously replicate in the genome, some TEs evolved or co-opted regulatory sequences to ensure their own transcription. As a consequence, the dispersed nature of TE regulatory sequences shapes the expression landscape of the genome (Mita and Boeke 2016; Chuong et al. 2017). Epigenetic silencing of the host genome and environmental triggers are major factors influencing TE transcription levels, although the underlying mechanisms are poorly understood.

Most TEs are transcriptionally and transpositionally quiescent (Yoder et al. 1997; Zilberman et al. 2007). However, environmental stimuli and stress, in particular, have been shown to trigger epigenetic de-repression of TEs resulting in the activation of insertional mutagenesis (Miousse et al. 2015). TE de-repression in response to stress is widely shared across eukaryotes (Bundo et al. 2014; Van Meter et al. 2014; Voronova et al. 2014; Romero-Soriano and Guerreiro 2016; Ryan et al. 2016; Zovoilis et al. 2016; Huang et al. 2017; Hummel et al. 2017; Shpyleva et al. 2018). De-repression of TEs under stress usually impacts TE transcription levels and can increase transpositional activity (Dubin et al. 2018). The impact of stress on TEs is often mediated through changes in the epigenetic state of the genome (*i.e.* de-repression) (Horváth et al. 2017) or the activation by a transcription factor (Capy et al. 2000). Some TEs have stress response elements (SRE) that are regulatory sequences activated in response to stress (Bucher et al. 2012; Casacuberta and González 2013). SREs are most common in long terminal repeat (LTR) retrotransposons and have been identified in one family of miniature inverted-repeat transposable elements (MITE) (Yasuda et al. 2013). The relationship between stress and TE activation is complex with some studies showing TEs being upregulated, some show TE repression and yet other studies show transient upregulation and then downregulation following exposure to a stress (Horváth et al. 2017). Stress mostly impacts facultative heterochromatin (Trojer and Reinberg 2007), while constitutive heterochromatin is typically associated with gene-poor, TE rich regions that maintain repression (Dillon 2004; Saksouk et al. 2015). The distribution of TE families or specific copies of a TE can be strongly correlated with the local chromatin state (Lanciano and Mirouze 2018). The epigenetic landscape influencing TE de-repression dynamics is a highly dynamic trait among closely related species (Niederhuth et al. 2016) but also showing significant variation within species (Barah et al. 2013).

TE responsiveness to stress potentially constitutes a major compound cost to the deleterious impact of stress on an organism. However, stress can induce both the activation and repression of TEs as was shown for different ecotypes of *A. thaliana* exposed to cold stress (Barah et al. 2013). In yeast and human cells TEs were found to be repressed in response to stress (Menees and Sandmeyer 1996; Trivedi et al. 2014). Another example in *Arabidopsis thaliana*, the *ONSEN* (LTR) retrotransposon is activated in response to heat stress due to heat response factors recognizing a regulatory sequence in the promoter of the ONSEN transposon (Ito et al. 2011; Cavrak et al. 2014). As a consequence, ONSEN insertions into genic regions were shown to induce the transcriptional upregulation of neighboring genes in response to heat stress (Ito et al. 2011). Therefore, TEs are frequently re-activated in response to stress and their activation can introduce new TE copies into the genome with *cis*-regulatory elements or associated chromatin states that are responsive to stress, thereby rewiring the stress response network of the genome (Cowley and Oakey 2013; Galindo-González et al. 2017). Hence, the stress activation of TEs likely depends on the type of stress, the identity of the TE and the genetic background of the host. Furthermore, TE activation may generate adaptive genetic variation and accelerate host stress adaptation.

TE de-repression dynamics in pathogens of plants show the hallmarks of a conflict between TE proliferation and host control. Insertions of TEs in pathogen genomes generate significant adaptive genetic variation through gene inactivation, gene copy-number variation and altered gene expression, and have been shown to play a role in the evolution of genes encoding proteins involved in host interaction (Croll and McDonald 2012; Seidl and Thomma 2017; Fouché et al. 2018). In fungi, TEs can also lead to genetic variation through repeat-induced point mutation (RIP), a genome defense mechanism that targets and mutates repetitive sequences (Selker 2002). In *Leptosphaeria maculans* for instance, leakage of RIP into neighboring regions contributes to the diversification of effector genes (Rouxel et al. 2011). During the infection of a host plant, the pathogen must overcome a number of severe stresses (Ferreira et al. 2006; Hernández-Chávez et al. 2017). Initially the pathogen is exposed to nutrient stress on the surface of the plant (Derridj 1996). Once the pathogen enters the plant, host defenses stimulate the accumulation of toxic reactive oxygen species (Shetty et al. 2007). To face plant-induced stresses and promote disease, pathogens express virulence factors (*i.e.* effectors). The expression of the virulence factors is often governed by de-repression of facultative heterochromatin (Connolly et al. 2013; Qutob et al. 2013; Chujo and Scott 2014; Soyer et al. 2014; Schotanus et al. 2015; Soyer et al. 2015; Studt et al. 2016). Infection stress incidentally serves as an epigenetic trigger for adaptive upregulation of effectors (Sánchez-Vallet et al. 2018). Concurrently, the pathogen genome becomes exposed to de-repressed TEs.

*Zymoseptoria tritici* is the most important pathogen of wheat in Europe (Fones and Gurr 2015; Torriani et al. 2015). The pathogen’s ability to infect host plants is largely determined by a complement of small proteins, most of them effectors, that manipulate the host physiology upon contact. Effector genes are frequently located in proximity to TEs and are highly up-regulated during early, stressful conditions of the host infection (Rudd et al. 2015; Palma-Guerrero et al. 2016; Haueisen et al. 2017; Palma-Guerrero et al. 2017; Fouché et al. 2018; Plissonneau et al. 2018). Effectors are thought to become upregulated by de-repression of facultative heterochromatin (Soyer et al. 2015; Soyer et al. 2019). Both facultative and obligate heterochromatin is highly enriched in TEs in *Z. tritici* (Schotanus et al. 2015). *Z. tritici* has a very plastic genome consisting of 13 core and up to eight accessory chromosomes that are not fixed within the species (Goodwin et al. 2011). Accessory chromosomes frequently undergo chromosomal rearrangements with breakpoints co-localized with TE insertions (Croll et al. 2013; Plissonneau et al. 2016; Hartmann et al. 2017; Plissonneau et al. 2018). Genes involved in pathogenicity and stress tolerance are frequently located in close proximity to TEs (Hartmann et al. 2017; Krishnan et al. 2018; Meile et al. 2018). For example, a TE insertion was shown to silence a transcription factor resulting in reduced melanin production (Krishnan et al. 2018). Furthermore, TE insertions contributed to the overexpression of a major facilitator superfamily (MFS1) transporter contributing to multi-drug resistance (Omrane et al. 2017). Populations segregate over a thousand gene presence-absence polymorphisms and gene deletions are preferentially located in proximity to TEs (Hartmann et al. 2017; Plissonneau et al. 2018).

In this study, we used transcriptome profiling to test for the impact of two major stress factors in the life-cycle of the pathogen on TE de-repression. We used a nutrient-rich culture medium as a non-stress environment and transferred the fungus to a nutrient-deprived medium that simulates starvation. Independently, we analyzed the fungal transcriptome at four distinct stages during the infection of wheat spanning the early symptomless stage, the peak of lesion formation and the saprotrophic stage. The early stages expose the pathogen to substantial nutrient and host defense stress factors. We replicated the two stress experiments with four genetically distinct strains of *Zymoseptoria tritici* differing in pathogenicity to identify how the genetic background influences TE responsiveness. TEs showed the highest expression under nutrient stress, but the expression differed significantly between TE families and between genetic backgrounds. Infection stress led to a large number of TE families to be upregulated at the peak of the symptom development on wheat leaves. Next, we determined how the genomic location affected the expression of TEs and identified distinct de-repression patterns depending on the type of stress, the distance to the closest genes and the impact of genomic defense mechanisms.

## Results

### TE landscape and transcriptomic response to stress conditions

We analyzed four strains of *Z. tritici* that differed significantly in the progression of infection and response to stress (Lendenmann et al. 2014; Lendenmann et al. 2016; Palma-Guerrero et al. 2017). The most virulent strain (3D7) developed visible symptoms within 12 days after infection (Palma-Guerrero et al. 2016). Strains 1A5 and 1E4 developed symptoms on average with a two-day delay and strain 3D1 showed the slowest symptom progression (Palma-Guerrero et al. 2017; Stewart et al. 2017). Each strain has a fully assembled and annotated genome (Plissonneau et al 2018) with similar percentages of TEs 16.0-18,1% (fig. 1A). Of the 111 TE families identified previously in the reference genome of the species (Grandaubert et al. 2015), all families were present in 1A5, 110 were identified in 1E4 and 3D1, and 108 were found in 3D7. LTR-*Gypsy* elements were the most abundant in all of the strains making up between 5-7% of the genomes, followed by LTR-*Copia* and LINE-*1* elements (fig. 1B). The TE content was highest in accessory chromosomes (14-21; fig. 1C). Chromosome 14 of strain 3D1 had the highest TE content (>40%). Despite the similarity in overall TE content between strains, TE superfamilies showed marked differences in their distribution across chromosomes (fig. 1C). SINE elements were only present on chromosome five for strain 1A5, 1E4 and 3D1 and on chromosome three for strain 3D7.

**Figure 1:**
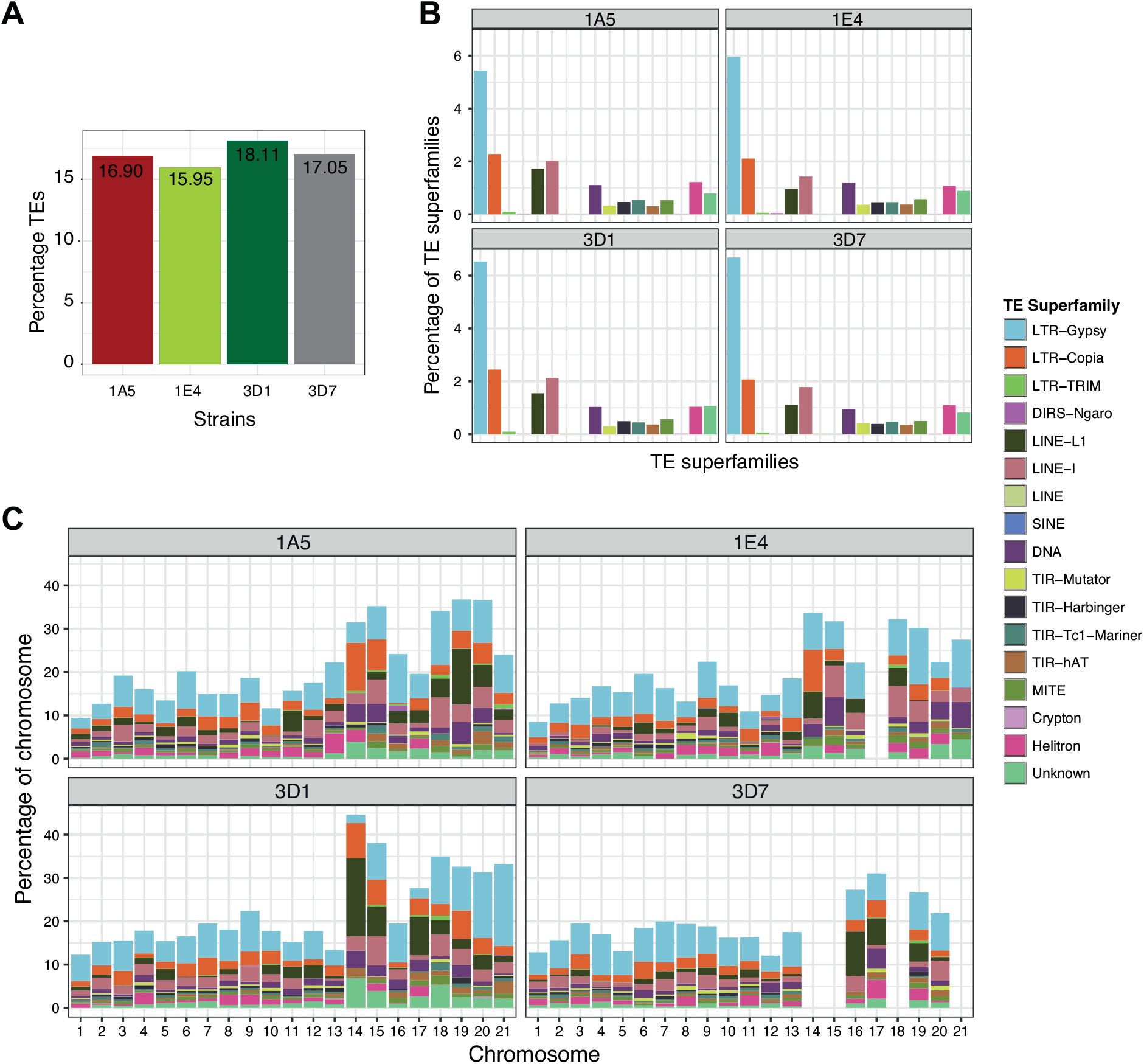
Transposable element (TE) composition of the completely assembled genomes of *Zymoseptoria tritici.* (A) The percentage TE in each genome. (B) The percentage of TE superfamilies in each strain. (C) The distribution of TE superfamilies as a percentage of the chromosomes for each strain. Although the percentage of TEs are similar between the strains, the distribution of elements along the chromosomes differ.

We analyzed the transcriptomic response to specific stress conditions by culturing the fungi first in nutrient-rich conditions, then analyzed the same strains growing in a minimal carbon source medium (*i.e.* starvation stress; fig. 2A). In parallel, we passaged all strains through an infection cycle on a wheat host (*i.e.* infection stress; fig. 2A). Infection stages were sampled at four time points (7, 12, 14 and 28 days post infection). Across all conditions, we found that biological replicates clustered tightly together showing high stress reproducibility (supplementary fig. S2). Gene expression profiles clustered mainly according to condition with early and late infection phases resembling nutrient starvation (fig. 2C). We analyzed the expression of putative virulence factors (*i.e.* effectors) and carbohydrate-active enzymes (CAZymes) in strain 3D7 in order to recapitulate the progression of the infection and impact of starvation. Overall, effector genes were upregulated during early infection stages (7-14 dpi) followed by downregulation at the final infection time point (supplementary fig. S3). CAZYmes are enzymes that digest carbohydrates and digest the plant cell walls, releasing nutrients for the pathogen. CAZYmes differed widely in expression profiles with subsets showing upregulation during early infection stages, in nutrient-rich conditions and nutrient starvation, respectively (supplementary fig. S4). Overall, stress conditions impose major gene expression profile changes consistent with the lifestyle transitions of the pathogen.

**Figure 2:**
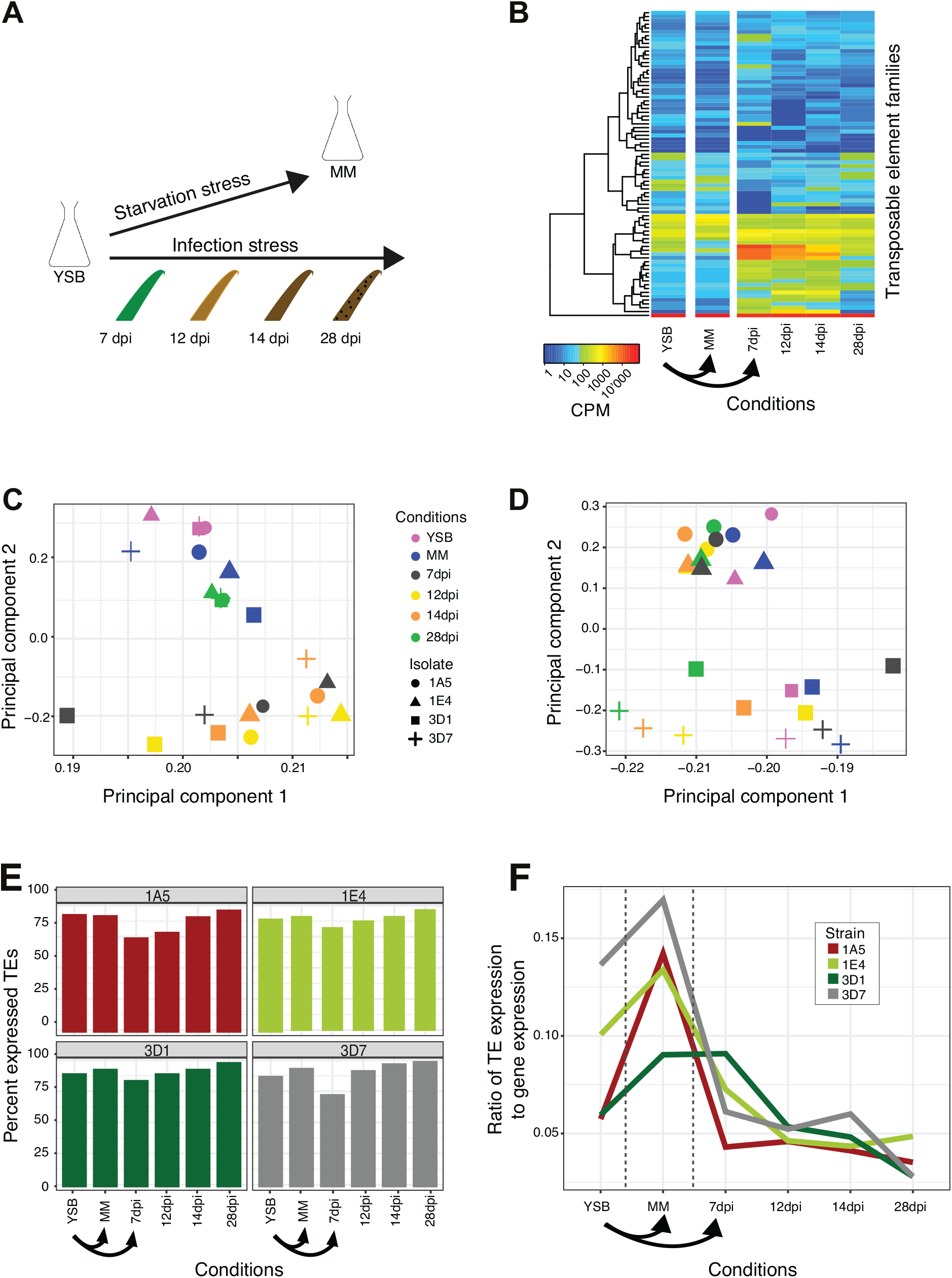
Responsiveness of transposable element (TE) to different stresses. (A) Transcriptomes analyzed in response to nutrient and infection stress. (B) Heatmap showing expression of TE families in the strain 3D1. Principal component analyses of gene expression (C) and TEs (D) in the four strains. (E) The percentage of expressed TEs compared to all TEs for each condition. (F) Proportion of TE expression to overall expression including genes.

### Differential stress response dynamics of TEs across environments

We analyzed TE expression across all conditions and strains using TETranscripts, which quantifies the expression abundance of a TE across all copies in the genome allowing for multiple read mapping. TE families differed substantially in expression profiles depending on the imposed stress condition (fig. 2B). A principal component analysis showed that the expression of TEs clustered according to the host genotype rather than stress condition (fig. 2D). Most TEs were expressed in most backgrounds and stress conditions (fig. 2E). The lowest percentage of expressed TEs was observed during early infection. We analyzed the relative expression of TEs versus genes expression and found that the highest relative TE expression occurred under nutrient-rich and starvation stress conditions (fig. 2F). The relative expression decreased with the progression of infection (fig. 2F).

The response of TEs to stress conditions was highly specific to individual TE families. In strain 3D7, two LTR-*Gypsy* element families and a TIR-*Tc1-mariner* element family, were only upregulated during early infection (7-14 days post infection; supplementary fig. S5B). Similarly, in strain 3D1 four LTR-*Gypsy* element families were mainly upregulated early during infection (supplementary fig. S5A). Two of these upregulated families were shared between the strains (LTR-*Gypsy* element families 6 and 9) and are the most infection stress responsive elements. In 1A5 and 1E4, a shared TE element family (TIR-*hAT* element 1) was most highly expressed during starvation and mostly repressed during infection (supplementary fig. S5C and D). Some TE families showed consistently high expression across all conditions suggesting generally weak genomic defenses against expression of this specific TE in comparison to other TEs that are only responsive to specific stress conditions (supplementary fig. S5A-D). TE expression in all four strains was dominated by a MITE-Undine family (fig. 3), which is the most highly upregulated TE under nutrient starvation stress in all strains. The exception is strain 3D7 where the family was similarly expressed under nutrient rich conditions and starvation stress (fig. 3A). MITE-Undine is a non-autonomous element lacking coding regions. We were unable to identify the helper autonomous element with the same terminal-inverted repeats (TIR). MITE-Undine was also the most abundant element in any of the four genomes (fig. 3B) with a copy number of 250-296. The mean number of TE copies per family in each genome was 29-32. The average distance of MITE-Undine to the nearest gene was 17.6-33.8 kb compared to 19.5-21.1 kb for TEs (supplementary table S2). The element was present on all chromosomes (fig. 3C for isolate 3D1) and contains target site sequences, TIR and low-complexity regions (palindromes and tandem repeats; fig. 3D). We found no evidence for the element in the genomes of the most closely related *Z. pseudotritici* and the more distantly related *Z. brevis*. However, *Z. ardabiliae* which has an intermediate divergence time from *Z. tritici* harbors 8 copies of the MITE-Undine.

**Figure 3:**
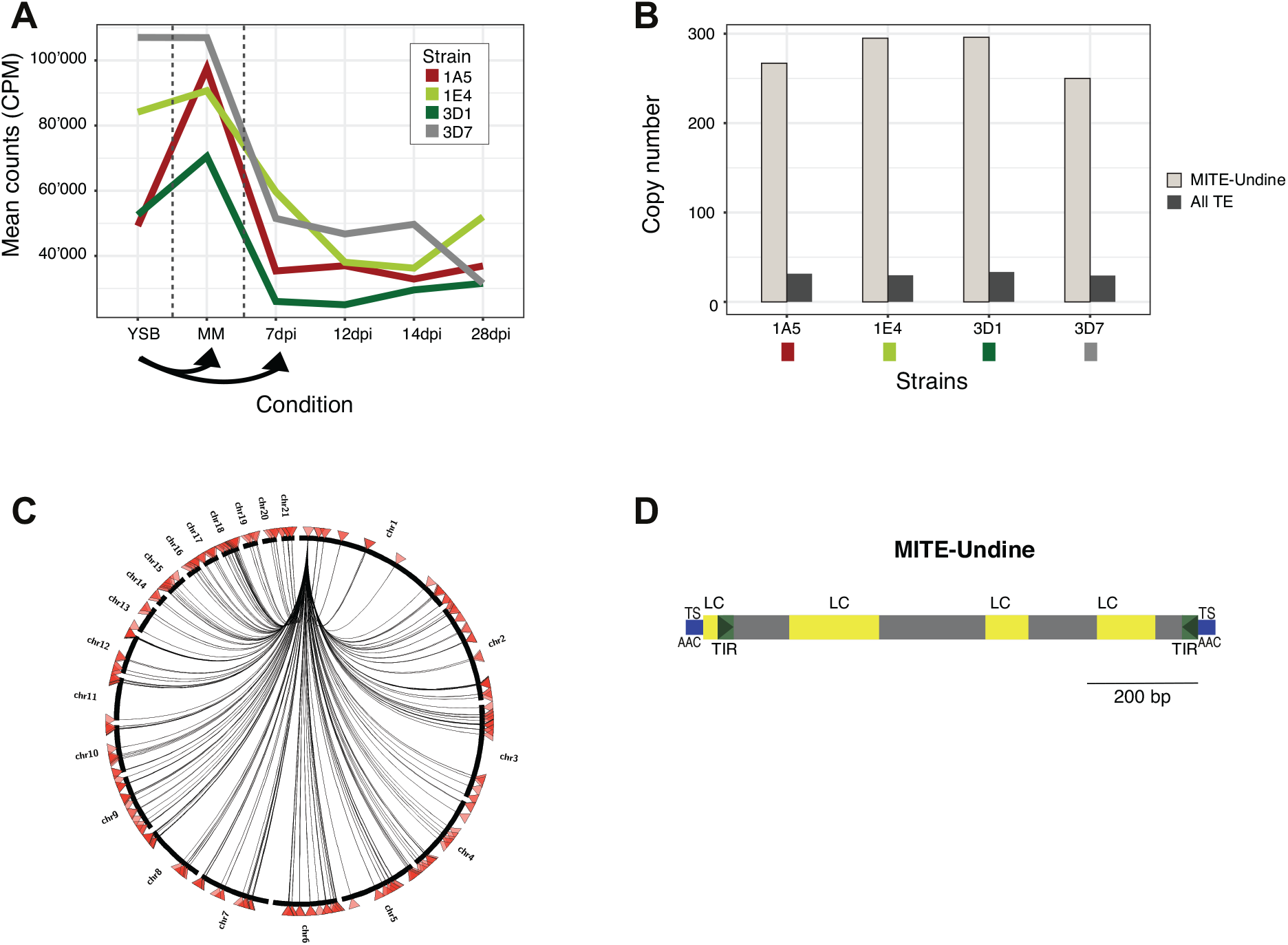
Characterization of the high-copy and highly-expressed MITE-Undine element. (A) The expression of MITE-Undine in counts per million (CPM) in all four strains. (B) The copy number of MITE-Undine compared to the mean copy number of all TEs. (C) Identification of MITE-Undine copies in the genome of 3D1. (D) Schematic of MITE-Undine with target site duplications (TS), terminal inverted repeats (TIR) and low complexity regions (LC).

### TE and gene expression dynamics as a function of the genomic localization

To address the impact of physical proximity to TEs on gene expression patterns, we analyzed gene expression across stress conditions as a function of distance to the closest TE. Genes within 1 kb upstream or downstream of TEs were upregulated early during infection (7-14 dpi depending on the strain) compared to genes >1 kb away from TEs (fig. 4A and supplementary fig. S6). Genes >1 kb away from TEs showed higher expression than genes <1 kb of TEs late in the infection (28 dpi). Genes with TE insertions showed consistently low levels of expression. Effector genes were overall closer to TEs than CAZYmes, genes encoding secreted proteins or genes overall (fig. 4B and supplementary fig. S7). Consistent with the proximity to TEs, effector genes were strongly upregulated early during infection compared to other gene categories (fig. 4C and supplementary fig. S8). Notably, the increase occurred first for 3D7, the strain with the most rapid infection progression (Palma-Guerrero et al. 2017) (supplementary fig. S8). The increase occurred at 12 dpi and peaked at 14 dpi for the other three strains (supplementary fig. S8).

**Figure 4:**
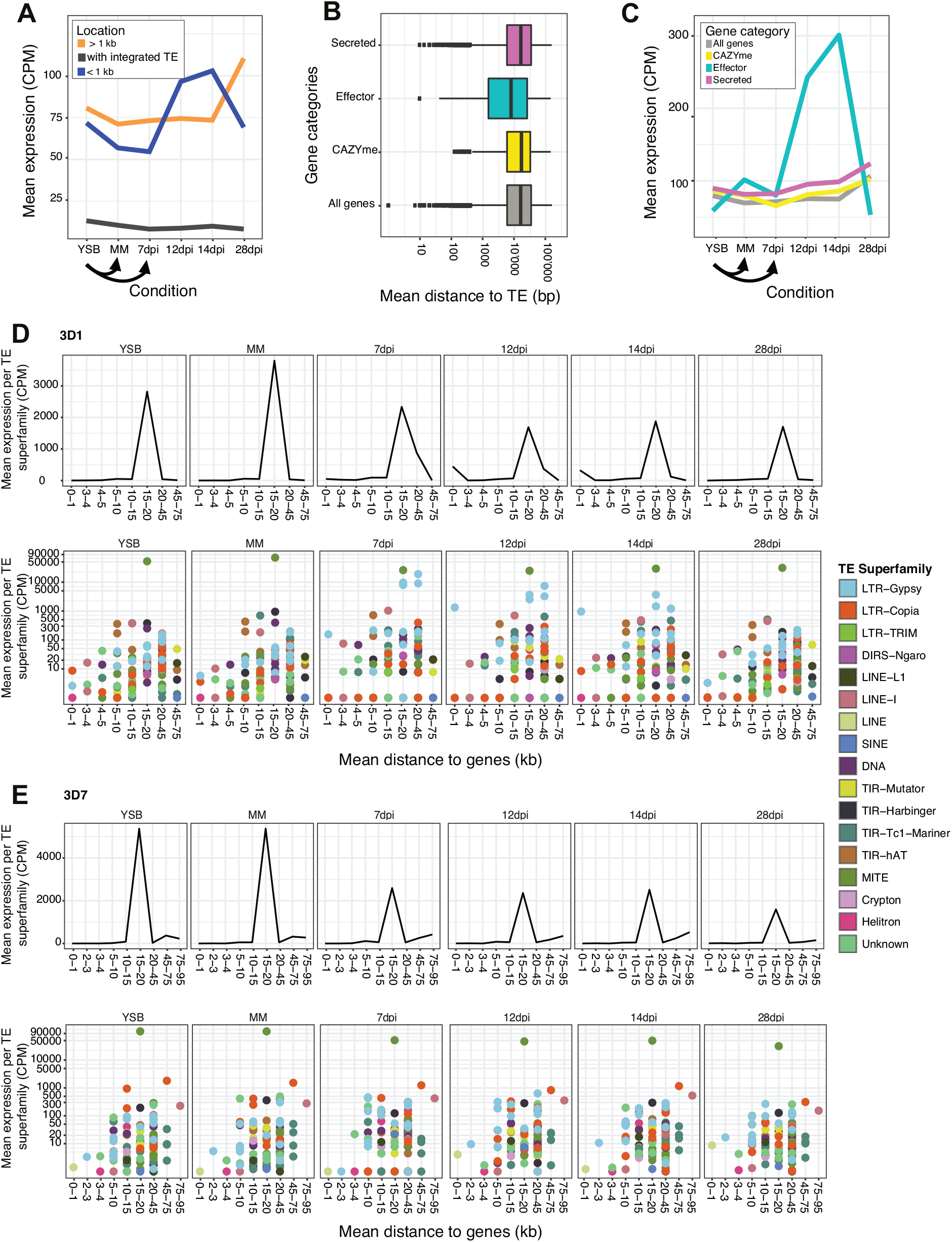
Gene expression as a function of proximity to transposable elements (TE) in strain 3D1. (A) Expression of genes with inserted TEs, within 1 kb of the nearest TE or more than 1 kb from the nearest TE. (B) Mean distance between genes of grouped by functional category (C) Mean expression of genes grouped by functional category. TE expression responses to stress as a function of the distance to the closest genes in 3D1 (D) and 3D7 (E). Distance segments lacking TEs were omitted.

Next, we analyzed TE expression responses to stress as a function of the distance to the closest genes. TE expression generally peaked for TEs at a mean distance of 15-20 kb to the closest gene for 3D1 and 3D7 and at 20-45kb away from the closest gene for 1A5 and 1E4 (fig. 4D and E and supplementary fig. S9). In strain 3D1 (Fig. 4D) and 1E4 (supplementary fig. S9B), TEs with a mean distance of within 1 kb of genes were upregulated during early infection. In 3D1, the upregulation at 7-14 dpi was primarily due to the expression of an LTR-*Gypsy* element, but also due to the expression of a LTR-*Copia* element family at 14 dpi (fig. 4D). In 1E4, TEs with a mean distance within 1 kb of genes were also upregulated under nutrient stress, primarily due to the expression of an unknown element and an LTR-*Gypsy* element (supplementary fig. S9B). In 3D7 and 1A5, TE families further away from genes were more strongly expressed (fig. 4E and supplementary fig. S9D). Exceptions in 1A5 include TE families with a mean distance of 1-2 kb to the nearest gene, which were upregulated under all of the conditions due to the expression of two unknown element families and two LTR-*Gypsy* families (supplementary fig. S9D). TE families with a mean distance of >45 kb away from the nearest gene showed upregulation under all conditions in strain 3D7, with a major peak at 12dpi (fig 4E). The most highly expressed families falling in this category were an LTR-*Copia* family at mean distance of 45-75 kb and MITE-Undine at 75-95 kb from the nearest gene.

### Co-expression networks of genes and TEs across stress conditions

Many TEs in the *Z. tritici* genome are in close physical proximity to genes and may, hence, converge on similar epigenetic expression dynamics across stress conditions. To infer synchronicity of TE and gene expression, we performed clustering analyses to define profiles of TE and gene co-regulation under stress. The analysis identified a total of twenty co-expression profiles, of which six were shared by all four strains and six were strain-specific. Eighteen co-expression profiles contained TEs, but were not identified in all isolates (fig. 5A). Expression profiles included different kinetics of upregulation upon infection stress (see profiles 16-17-18-27-28-29-31-39) but also downregulation (see profiles 8-9-13-15-19-20-21-32-35) with various intermediary profiles (fig. 5A). Co-expression profiles included on average >98% of genes and up to 5% of TEs in each genome (fig. 5A). To infer the biological relevance of different co-expression clusters, we performed enrichment analyses of gene ontology (GO) terms. In total, 11, 7, 14 and 12 co-expression profiles showed significant enrichment for GO terms in strains 1A5, 1E4, 3D1 and 3D7, respectively (p-value < 0.05). Four co-expression profiles were found consistently enriched for GO terms in the four strains (profiles 8, 18, 32 and 36; fig. 5A). Profile 8 was >2-fold enriched for phosphorylation activity and transcriptional activity among the four strains (supplementary table S3). Profile 18 showed consistent enrichment for GOs related to carbohydrate metabolism, hydrolase activity, proteolysis and ribosome (up to 15-fold; supplementary table S3). Profile 32 was enriched for genes encoding kinases, protein transporters and DNA replication functions. Profile 36 was enriched for metabolic processes, proteolysis and catalytic activity (supplementary table S3). In the strains 1A5, 1E4 and 3D7 the co-expression profile 39 showed a consistent upregulation from starvation stress (MM) to late infection and was enriched (up to 20-fold) for GO terms related to ribosome and ATP synthesis (fig. 5a; supplementary table S3).

**Figure 5:**
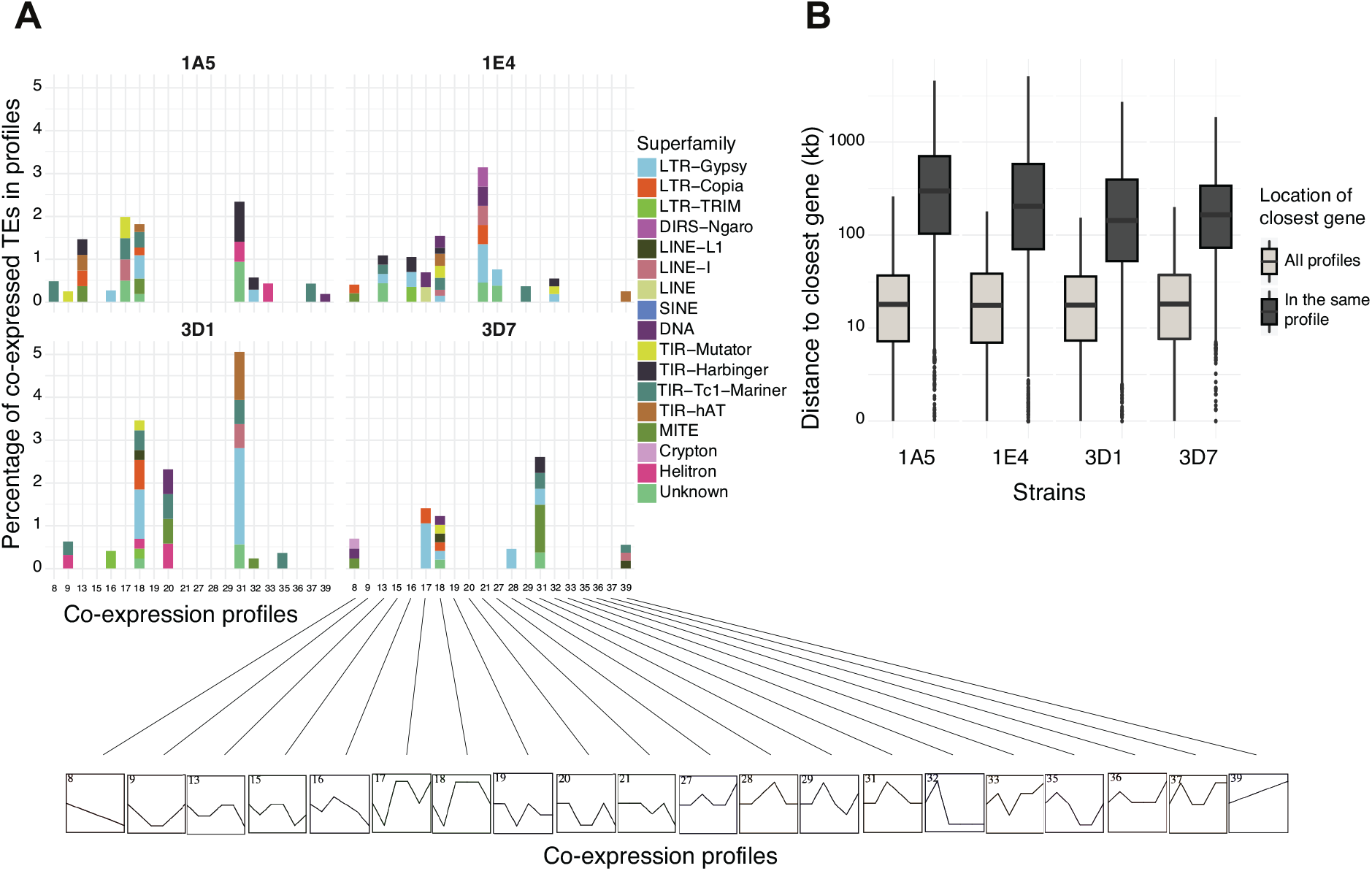
Co-expression profiles of transposable elements (TE) and genes. (A) The percentage of TE families in each co-expression profile compared to the total number of. (B) The distribution between TEs and co-expressed genes. TEs and co-expressed genes are not closer than TEs and genes that do not share an expression profile.

Both DNA and retrotransposons were co-expressed with genes. LTR-*Gypsy* elements were dominating co-expression profiles reflecting the abundance of the elements in the genomes. The number of co-expressed TE superfamilies ranged from 1 to 8 per profile. Among all four strains, the co-expression profiles displaying higher numbers of TE superfamilies were consistently those with a peak of expression early in the infection process (*e.g*. profile 18). Overall these results reveal that multiple TEs and diverse superfamilies are found co-expressed together with genes upon host infection. TE and gene co-regulation could be driven by shared epigenetic environments due to physical proximity and/or transcriptional leakage. To test for the effect of physical proximity, we analyzed the physical distance between co-expressed TEs and their closest co-expressed genes. In concordance with the previous global analysis, co-expressed genes with a peak of expression early upon infection are found closer to TEs. However, the closest distance between co-expressed genes and TEs within an expression profile is on average ten times longer than the distance of the closest co-expressed genes and TEs not in the same expression profile (fig. 5B). Therefore, TEs and co-expressed genes are not closer than TEs and genes that do not share an expression profile.

### Impact of genomic defenses on TE expression under stress

Fungi evolved sophisticated genomic defenses that inactivate TE copies through the introduction of RIP mutations (Selker 2002). In order to determine how RIP may impact TE expression under stress, we analyzed mutational biases among genomic TE copies. Most TE families in all four genomes were affected by RIP (fig. 6A and supplementary fig. S11). Only TE families in the LTR-TRIM superfamily and one family in the TIR-*Tc1*-*mariner* superfamily were not affected by RIP. In all strains, TEs in the TRIM family were among the most highly affected TEs. In 1A5, a family belonging to the TIR-*Tc1-mariner* superfamily is consistently expressed under all conditions (supplementary fig. S10A). Similarly, LTR-*Gypsy* elements with strong RIP signatures were upregulated upon infection in the strains 1A5, 1E4 and 3D1 (supplementary fig. S11A). Hence, most TE families affected by RIP are still expressed under at least some stress conditions.

**Figure 6:**
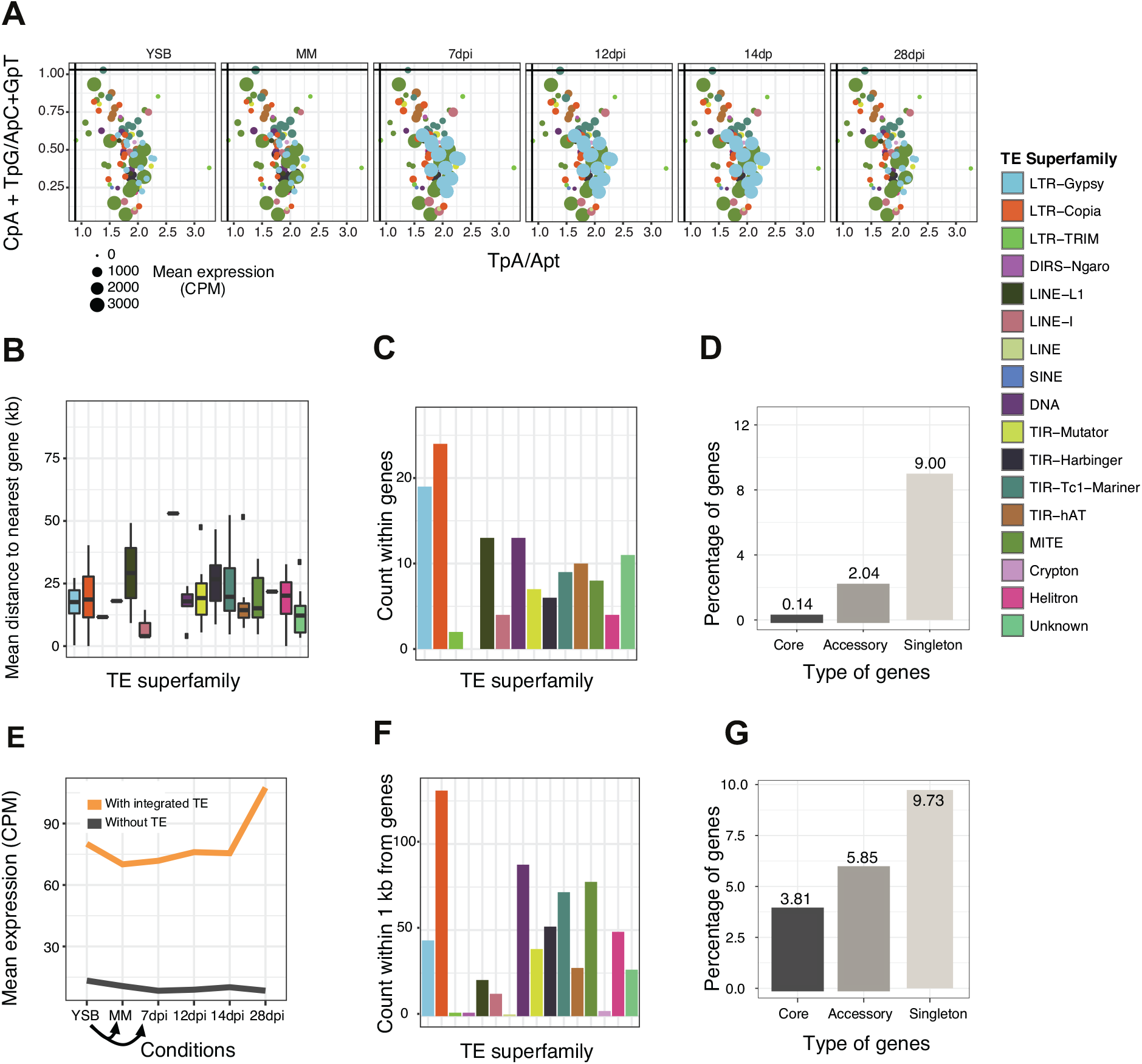
Genomic defenses and transposable element (TE) expression. (A) Repeat induced point mutation (RIP) indices for each TE family and mean expression of the family under all stress conditions for strain 3D1. Vertical and horizontal lines represent commonly used thresholds to detect RIP (Hane and Oliver 2008). Colours indicate the superfamily and size the expression at the family level in CPM. (B) The mean interval of TE superfamilies to the nearest gene. (C) TE superfamilies within genes in 3D1. (D) Gene categories with inserted TEs in 3D1. The percentages are given as the number of genes of that category (core accessory or strain specific) within genes as a fraction of the total number of genes in a category. (E) Expression of genes with or without inserted TEs. (F) TE superfamilies within 1kb of the closest gene in strain 3D1. (G) Gene categories within 1 kb of TEs. The percentages are given as the number of genes of that category (core accessory or strain specific) within genes as a fraction of the total number of genes in a category.

### TE insertion dynamics in proximity to genes

TE superfamilies showed substantial variation in their mean distance to the closest gene with most having a mean distance to the nearest gene of less than 25 kb (fig. 6B and supplementary fig S12). The closest TE superfamilies to genes were DIRS-Ngaro (2867 bp) in 1A5, LINE (415 bp) in 1E4, LINE-*1* (7457 bp) in 3D1 and LINE (832 bp) in 3D7 (supplementary table S4). Next, we analyzed coding sequence disruptions across the genome and found 169, 136, 130 and 103 TEs inserted into genes in strains 1A5, 1E4, 3D1 and 3D7, respectively. LTR-*Copia* elements were the most frequently inserted TEs into genes in strains 1E4, 3D1 and 3D7 (22, 24 and 26 insertions, respectively), while in 1A5 LINE-*L1* elements were most often (27 insertions) inserted into genes followed by elements belonging to the LTR-*Copia* superfamily (25 insertions; fig. 6C and supplementary fig. S12A and supplementary table S5). Singleton genes defined as present in only one of the four strains were the most frequently disrupted genes (fig. 6D and supplementary fig. S12B). Genes with inserted TEs had a lower expression than genes without TEs (fig. 6E and supplementary fig. S12E). Hereafter, we analysed TE insertions in close proximity to genes. We found 605-683 TEs <1 kb from genes among strains (excluding within gene insertions) and LTR-*Copia* elements were the most abundant elements with 108-132 copies (fig. 6F and supplementary fig. S12C and supplementary table S6). Singleton genes most frequently had an integrated TE or were located within 1 kb from a TE, followed by accessory genes and core genes (fig. 6D and G and supplementary Figure S12B and D).

## Discussion

TEs are major drivers of genome evolution due to their transpositional activity. Repression of TEs is largely governed through epigenetic control and is, hence, susceptible to external stress. Using transcriptome profiling, we show that two distinct stress conditions induce the expression of distinct sets of TEs. By replicating the analyses across four genetic backgrounds, we show that the major expression dynamics of TEs are conserved. However, some of the most highly expressed TEs including MITE and LTR-*Gypsy* elements showed highly distinct de-repression across stress conditions. The genomic context of TEs was a major predictor of de-repression dynamics during stress. Consistent with TE de-repression being governed by epigenetic effects, we found that gene expression profiles under stress varied significantly depending on the proximity to the closest TEs. Genomic defenses capable of introducing point mutations into repetitive sequences (*i.e.* RIP) affected most TE families, however a large subset of affected TEs were still de-repressed under stress.

### Stress-dependent TE de-repression dynamics

The completely assembled genomes of *Z. tritici* display substantial variability in chromosome-level TE content despite highly similar overall repetitive element proportions. The TE content variation is striking given the fact that all four strains were collected from nearby wheat fields, interfertile and from populations with a rapid decay in linkage disequilibrium (Croll et al. 2015). LTR-*Gypsy* were the most abundant elements consistent with their abundance in many other fungal genomes (Muszewska et al. 2011). Members of the LTR-*Gypsy* superfamily in conjunction with a MITE showed among the strongest de-repression under stress. MITEs and LTR-retrotransposons are also most frequently associated with stress responsiveness in other organisms (Yasuda et al. 2013; Negi et al. 2016). However, the impact of stress on TE expression is highly variable among TE families, copies and species. Some TEs are expressed and potentially mobilized in response to stress, while other TEs are suppressed after an initial stress-induced activation, and some TEs are downregulated in response to stress (Horváth et al. 2017).

Nutrient starvation and host infection constitute the major stress factors in the life cycle of filamentous plant pathogens (Ferreira et al. 2006; Hernández-Chávez et al. 2017). We exposed *Z. tritici* to two stress conditions. Growth in a carbon source depleted culture medium (MM) exposed the fungus to nutrient starvation. Early infection stages induce stress due to host immune responses targeted at the pathogen and imposes growth under limited nutrient conditions. Carbohydrate-active enzymes (CAZYmes) showed highly distinct profiles depending on the stress condition. Hence, starvation and infection stress have distinct impacts on gene expression consistent with the biological context of the stress (Palma-Guerrero et al. 2016). The majority of TE families showed some degree of de-repression in at least one of the stress conditions. Elements in the LTR-*Gyps*y superfamily were upregulated during early infection, which corresponds to the most stressful period on the host (Rudd et al. 2015; Palma-Guerrero et al. 2016). Infection stress first causes the upregulation of effector genes and later cell-wall degrading enzymes (Skibbe et al. 2010; Wang et al. 2011; Kleemann et al. 2012; Hacquard et al. 2013; Soyer et al. 2014; Palma-Guerrero et al. 2016). Effector gene expression is known to be epigenetically regulated in plant pathogens and timed to maximize exploitation of the host (Qutob et al. 2013; Schotanus et al. 2015; Sánchez-Vallet et al. 2018; Soyer et al. 2019). We found that a different set of TEs showed the highest expression under starvation stress including a MITE, which was the most strongly expressed TE in the genomes. The regulatory framework governing stress responses is largely unknown in *Z. tritici*, but distinct epigenetic regulation in response to stress is likely playing a key role (Schotanus et al. 2015; Soyer et al. 2019).

### TE and gene co-expression dependent on the genomic environment

Genes and TEs in close physical proximity likely undergo joint epigenetic regulation in response to stress. We found that genes close to TEs were upregulated during early infection consistent with the de-repression observed for TEs. LTR-*Copia* elements were the most frequently found TEs close to genes and one LTR-*Copia* family showed upregulation during early infection. In contrast, genes far from TEs were upregulated towards the end of the infection cycle, which is after the transition to a less stressful saprophytic lifestyle (Palma-Guerrero et al. 2016). Interestingly, we found no association of gene-to-TE distance with expression under nutrient starvation stress. This distinction may be due to the fact that epigenetic control of TEs is less pronounced under nutrient starvation stress. To understand TE de-repression dynamics as a function of the genomic environment, we also analyzed mean distances of TE families to the closest genes. Due to the repetitive nature of TEs, most transcriptome-derived short sequences cannot reliably be assigned to a single TE copy. Hence, our distance analyses were performed using summary statistics per TE family and not per individual TE copy. Copies of the most highly expressed TE, a MITE-Undine, are 17.6-33.8 kb away from genes across all genetic backgrounds. This activation could still be affecting the expression of genes as was shown for the Hopscotch TE in maize. This TE influences the expression of the TB1 locus at a distance of ~60kb (Studer et al. 2011).

Based on our co-expression clustering analyses, we found that TEs were not physically closer to co-expressed genes than other genes, suggesting that co-regulation is occurring in *trans* rather than in *cis*. Alternatively, this may reflect the epigenetic landscape of the genome with a multitude of distal chromosomal regions showing concerted de-repression dynamics. In other organisms, the influence of a TE on nearby genes is largely determined by the chromatin state of the TE (Saze and Kakutani 2007; Martin et al. 2009; Zeng and Cheng 2014; Lei et al. 2015; Williams et al. 2015; Williams et al. 2015; Hirsch and Springer 2017). This is most evident for stress responsive genes that carry TE insertions in the promoter sequences leading to upregulation upon TE demethylation (Le et al. 2014). Whether TE silencing through chromatin modification can spread to adjacent genes is not well understood (Sienski et al. 2012; Le Thomas et al. 2013; Lee 2015). TE stress responsiveness can be governed by epigenetic de-repression (Slotkin and Martienssen 2007) or the loss of a repressive mechanism under stress (Van Meter et al. 2014). If all TEs showed a correlated response to stress, this would suggest that activation is mostly due to epigenetic effects alone. However, different TE families show different stress responsiveness according to the stress condition and the host genotype, suggesting distinct epigenetic environments between the strains.

### The role of the genetic background in TE expression dynamics

The set of four completely assembled con-specific genomes enabled us to analyze de-repression dynamics of the same TEs across different genetic backgrounds. We found that TE family expression differed between the strains indicating that the genetic background plays a role in the ability of TEs to respond to stress. Several LTR-*Gypsy* elements were upregulated early during infection in one genetic background but not in all. TE families in close proximity to genes were upregulated during early infection in strains 3D1 and 1E4, while families with the longest average distance to genes were upregulated during infection in strain 3D7. Our evidence for TE family expression by genetic background interactions suggests high degrees of polymorphism for TE control within the species. In *A. thaliana*, TE responsiveness to cold stress was found to differ among ecotypes and this was largely explained by differences in the genomic locations of specific TEs (Barah et al. 2013). Such variation in the ability to control TE expression provides selectable genetic variation for the host genome to evolve more efficient control mechanisms.

### Stress de-repression of TEs and the evolution of TE control mechanisms

De-repressed TEs are mutagenic (Le Rouzic and Capy 2005) and can lead to genome expansions (Lonnig and Saedler 2002). Hence, host genomes evolved to suppress TE proliferation. Stress responsiveness of TE families is indicative of the ability by the host to control proliferation. The MITE with the highest expression is consistently expressed in all conditions, suggesting that the host genome has not yet evolved effective control mechanisms. RIP is a genomic defense mechanism that hypermutates duplicated DNA sequences in fungi and counteracts TE proliferation (Selker 2002). We found that TE families with signatures of RIP were still responsive to stress. In particular, the highly responsive MITE and LTR-*Gypsy* elements under starvation and infection stress, respectively, display strong signatures of RIP. This suggests that point mutations introduced by RIP may well introduce loss-of-function mutations and disable *e.g.* transposase functions. However, RIP in *Z. tritici* seems ineffective at preventing TE de-repression under stress.

Pathogens of plants are exposed to unique stress conditions upon entering their host. The challenges mounted by the plant immune system are designed to effectively contain a pathogen’s deployment of its infection program. Specialized pathogens evolved to time the expression of pathogenicity factors with the onset of stress by localizing the underlying genes in epigenetically silenced chromosomal regions (Sánchez-Vallet et al. 2018). We show here that the co-localization of epigenetically silenced TEs and genes leads to a highly correlated expression response upon infection. While the localization of pathogenicity factors in epigenetically silenced regions is most likely adaptive for the pathogen, the localization of TEs in the same compartment is likely only adaptive in absence of stress. Hence, the co-localization of pathogenicity factors and TEs creates a complex selection regime on the pathogen. Selection for more effective TE control under infection stress may actually be deleterious for the coordinated gene expression during infection. We identified unexpected complexity in both the genomic localization of TEs across genetic backgrounds and in the TEs response to stress. This suggests that there is standing variation for the ability to control TEs within the species. Hence, host genomes and TEs may be engaged in rapid co-evolutionary arms races to maintain effective control and escape repression, respectively.

## Methods and materials

### Strains and growth conditions

Strains 1A5, 1E4, 3D1 and 3D7 were isolated from two fields in Switzerland in 1999 and have been phenotypically and genotypically characterized (Zhan et al. 2005; Croll et al. 2013). These strains were used in previous studies for quantitative loci mapping (Lendenmann et al. 2014; Stewart and McDonald 2014; Lendenmann et al. 2016; Stewart et al. 2017). The genomes of all four strains have been sequenced and assembled into complete chromosomes using high-coverage PacBio sequencing (Plissonneau et al. 2016; Plissonneau et al. 2018). High-density genetic maps (Lendenmann et al. 2014; Croll et al. 2015) were used to validate each assembly.

### Gene and repetitive element annotation

We used pangenome gene annotation generated by Plissonneau et al. (2018). Genes were predicted by using splicing evidence from the *in planta* RNA-seq data from the same time points and strains as described above. Repetitive elements were annotated in all four genomes using RepeatMasker 4.0.5 (Smith, 1996) and a repeat element library for the reference genome (IPO323) produced by Grandaubert et al. (2015). This library was created using the REPET pipeline (Flutre et al. 2011). Repeat families in each species were clustered with BLASTClust from the NCBI-BLAST package (Altschul et al. 1990) and aligned with MAFFT (Katoh and Standley 2013) to create new consensus sequences. This process was repeated with lower identity percentages (from 100% to 75% identity) and lower coverage (from 100% to 30%) until sequences did not form new clusters anymore. The repetitive sequences were classified with the TEClassifier.py script from REPET using tBLASTx and BLASTx against the Giri Repbase Update database (Jurka et al. 2005) and by the identification of characteristic TE features such as LTRs. The sequences were also translated into the six reading frames in order to identify protein domains in the conserved domain database (Marchler-Bauer et al. 2011) using RPS-BLAST. Identified repetitive sequences were finally named according to the three-letter nomenclature defined by Wicker et al. (2007). The single TE library enables the comparison of TEs between the four strains as all the elements have exactly the same naming. RepeatMasker was used with the following parameters: pa 2, -s and –a for using two parallel processors, in slow mode for increased sensitivity and generating an alignment output file. Additional elements were identified using MITEtracker with default parameters (Crescente et al. 2018).

### RNA extractions, library preparation and sequencing

Seedlings of the wheat cultivar ‘Drifter’ were infected with the four strains 3D1, 3D7, 1E4 and 1A5, on the same day and in the same greenhouse chamber (Palma-Guerrero et al. 2016; Palma-Guerrero et al. 2017). Total RNA was extracted from inoculated second leaves at time points 7, 12, 14 and 28 days post infection (dpi) using a TRIzol (Invitrogen) extraction protocol (Palma-Guerrero et al. 2017). The time points were selected to include asymptomatic (biotrophic), necrotrophic and the saprophytic stages of infection. In addition, a TRIzol RNA extraction was performed for all four strains in a nutrient limited, defined salts medium without sucrose (Minimal Medium – MM pH 5.8) and a nutrient rich YSB media (10g/L sucrose and 10g/L yeast extract, pH 6.8) (Vogel et al. 1956; Francisco et al. 2018). Cells were recovered in YSB medium and then transferred to either YSB or MM, incubated for four days at 18°C, prior to harvest for RNA extraction. For the infection experiments, cells were harvested from three leaf samples from each time point for the *in planta* samples and three *in vitro* samples for each condition. Samples with the highest RNA quality as determined with a Bioanalyzer 2100 (Agilent), were selected as biological replicates for each time point for library preparation and sequencing. RNA quantity was assessed with a Qubit fluorometer (Life Technologies) and libraries were prepared using the TruSeq stranded mRNA sample prep kit (Illumina Inc.) according to the provided protocol. Total RNA samples were ribosome depleted by using PolyA selection and reverse-transcribed into double-stranded cDNA. Actinomycin was added during the first strand synthesis. The cDNA was then fragmented, end-paired and a A-tail was added before the ligation of the TruSeq adapters. A selective enrichment for fragments with TruSeq adapters on each end was performed by polymerase chain reaction. The quality and quantity of the enriched libraries were verified with a Qubit (1.0) fluorometer and a Tapestation (Agilent). Paired-end libraries were sequenced on an Illumina HiSeq 2500 with read lengths of 2 × 125 bp (Illumina Inc.) for *the in planta* samples and with read lengths of 4 × 100 bp for the *in vitro* conditions.

### Transcription mapping and quantification

Raw sequencing reads were quality-trimmed and filtered for adapter contamination and low-quality reads using Trimmomatic 0.36 (Bolger et al. 2014) using the following parameters: ILLUMINACLIP:TruSeq3-PE.fa:2:30:10 LEADING:10 TRAILING:10 SLIDINGWINDOW:5:10 MINLEN:50. Trimmed and filtered reads were mapped to the reference genome sequence of the specific strain (Plissonneau et al. 2018) using STAR 2.6.0 (Dobin and Gingeras 2016) allowing multiple mapped reads with the following settings: --outFilterMultimapNmax 100 --winAnchorMultimapNmax 200 --outSAMtype BAM Unsorted --outFilterMismatchNmax 3, according to the recommended parameters for TE analyses (Jin et al. 2015; Jin and Hammell 2018). We performed a saturation analysis to determine the cut-off to be used for the optimal number of reported alignments of a specific read, where increasing the threshold did not increase the number of mapped reads significantly as recommended (Jin and Hammell 2018) (supplementary fig. S1, supplementary table S1). The resulting unsorted bamfiles were sorted by read name with SAMtools 1.9 and the expression levels of TEs and genes were quantified using TEtranscripts 2.0.3 (Jin et al. 2015) with the following parameters: --stranded no --mode multi -p 0.05 -i 10. TEtranscripts counts uniquely and multiple-mapped reads that align to genes and TE regions to determine TE- and gene-level transcript abundance. The software assumes that transcribed TEs will have reads mapping along the entire length of the element (Jin and Hammell 2018). Elements with reads mapping to only a fraction of the length were assumed to be non-transcribed as these subregions may not be unique enough in the genome compared to *e.g.* other TEs.

TE and gene read counts were normalized between replicates and time points for the *in planta* and *in vitro* samples using the R/Bioconductor package EdgeR 3.8 (Robinson and Smyth 2008; Robinson et al. 2010; Robinson and Oshlack 2010). Genes and TEs without at least one read in all the samples were excluded for the normalization step (Anders et al. 2013) and were assumed for the rest of the analyses to have zero expression. Library sizes were normalized with the TMM method (Robinson and Oshlack 2010). CPM (counts per million) were generated with the EdgeR CPM function using TMM-normalized libraries.

### Genomic localization of TEs and co-expression analyses

In order to investigate the association of the genomic environment with TE expression, we identified the nearest gene to each TE using bedtools 2.27 command *closest* with the option to report only the closest TE to each gene and to allow overlaps to include genes that have been disrupted by intragenic TEs (Quinlan and Hall 2010). Co-expression clusters were computed using the Short Time-Series Expression Miner (STEM) software 1.3.11, designed to analyze time series with 3-8 time points (Ernst et al. 2005; Ernst and Bar-Joseph 2006). STEM software uses a non-parametric clustering method to assign genes to predefined expression profiles. It considers expression profiles to be significant if the number of genes assigned to a cluster departs from random. We used all three individual replicates per condition and isolate (option: repeat data) and transformed all data using log normalization. The analysis identified a total of 20 co-expression profiles, namely 8-9-13-15-16-17-18-19-20-21-27-28-29-31-32-33-35-36-37 and 39. The statistical significance of the number of genes assigned to each profile was computed by applying a Bonferroni correction with alpha = 0.05. The biological relevance of co-expression profiles was assessed by gene ontology (GO) term enrichment analysis (Ernst et al. 2005; Ernst and Bar-Joseph 2006). STEM software implements a GO term enrichment method that uses the hypergeometric distribution based on the number of genes assigned to the co-expression profile, the number of genes assigned to the GO category and the number of unique genes in the experiment. Enrichment significance was corrected by using randomization tests.

### RIP analysis

We identified RIP by aligning each copy of a given TE with the consensus sequence using MAFFT 7.407 (Katoh et al. 2002; Katoh and Standley 2013). The consensus sequence could be more affected by RIP than individual TE copies in the genome because mutations occurring in a given sequence are likely to be removed by the “base-pair majority rule” used to build the consensus. In this case, the copy with the highest GC content (*i.e*. the least affected by RIP) is used as the RIPCAL 1.0 input (Hane and Oliver 2008). All TE families (individual TE copies and the consensus) were aligned and processed by RIPCAL using default parameters (Hane and Oliver 2008). RIPCAL output provides the number of transition and transversions, single mutations and dinucleotide targets used in all possible transition mutations for each genomic TE copy. The RIPCAL output can be used to determine whether individual TE copies are “RIPped” based on two indices: (CpA+TpG)/(ApC+GpT) indicating a decrease in RIP targets and TpA/ApT, indicating an increase in RIP products. We used the default criteria where a (CpA+TpG)/(ApC+GpT) ratio of below 1.03 is indicative of RIP and a TpA/ApT of higher than 0.89 is indicative of RIP (Hane and Oliver 2008). In general TE families with a lower (CpA+TpG)/(ApC+GpT) value and a higher TpA/ApT are more affected by RIP (Hane and Oliver 2008). We excluded unknown elements from this analysis. R (R Core Team, 2017) was used to generate graphics from RIPCAL outputs. These outputs were parsed to search for RIP signatures in TE copies and the dinucleotide targets used in the transition type mutations that are usually associated with RIP.

## Data availability

The genome assembly and annotation for 1A5, 1E4, 3D1 and 3D7 genome are available at the European Nucleotide Archive (http://www.ebi.ac.uk/ena) under accession numbers PRJEB15648, PRJEB20900, and PRJEB20899 and PRJEB14341.The *in plant*a RNA-sequencing raw datasets are available at the NCBI Short Read Archive under accession number SRP077418. The *in vitro* RNA-sequencing raw sequencing data were deposited into the NCBI Short Read Archive under the accession number SRP152081.

## Author contributions

Conceived the study: SF, CP, DC Performed analyses: SF, TB, UO, CP Contributed dataset: CSF Wrote the manuscript: SF, DC

## Acknowledgements

We thank Javier Palma-Guerrero for providing access to transcriptomic datasets and Emilie Chanclud for helpful comments on a previous version of the manuscript. Data produced in this paper were generated in collaboration with the Genetic Diversity Centre (GDC), ETH Zurich and the Functional Genomics Center Zurich. SF was supported by the Swiss National Science Foundation (http://www.snf.ch) through the grant (SNF 31003A_155955) awarded to Bruce A. McDonald. DC receives support from the Swiss National Science Foundation (grants 31003A_173265 and IZCOZO_177052).

