## Supplementary Information for "Stress-driven transposable element de-repression dynamics in a fungal pathogen"

### Supplementary Figures

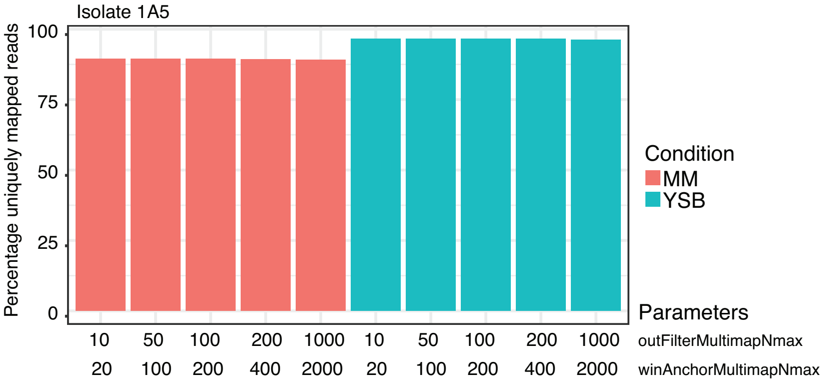


**Supplementary Figure S1: Saturation analysis to determine the cut-off to be used for the optimal number of reported alignments of a specific read.**

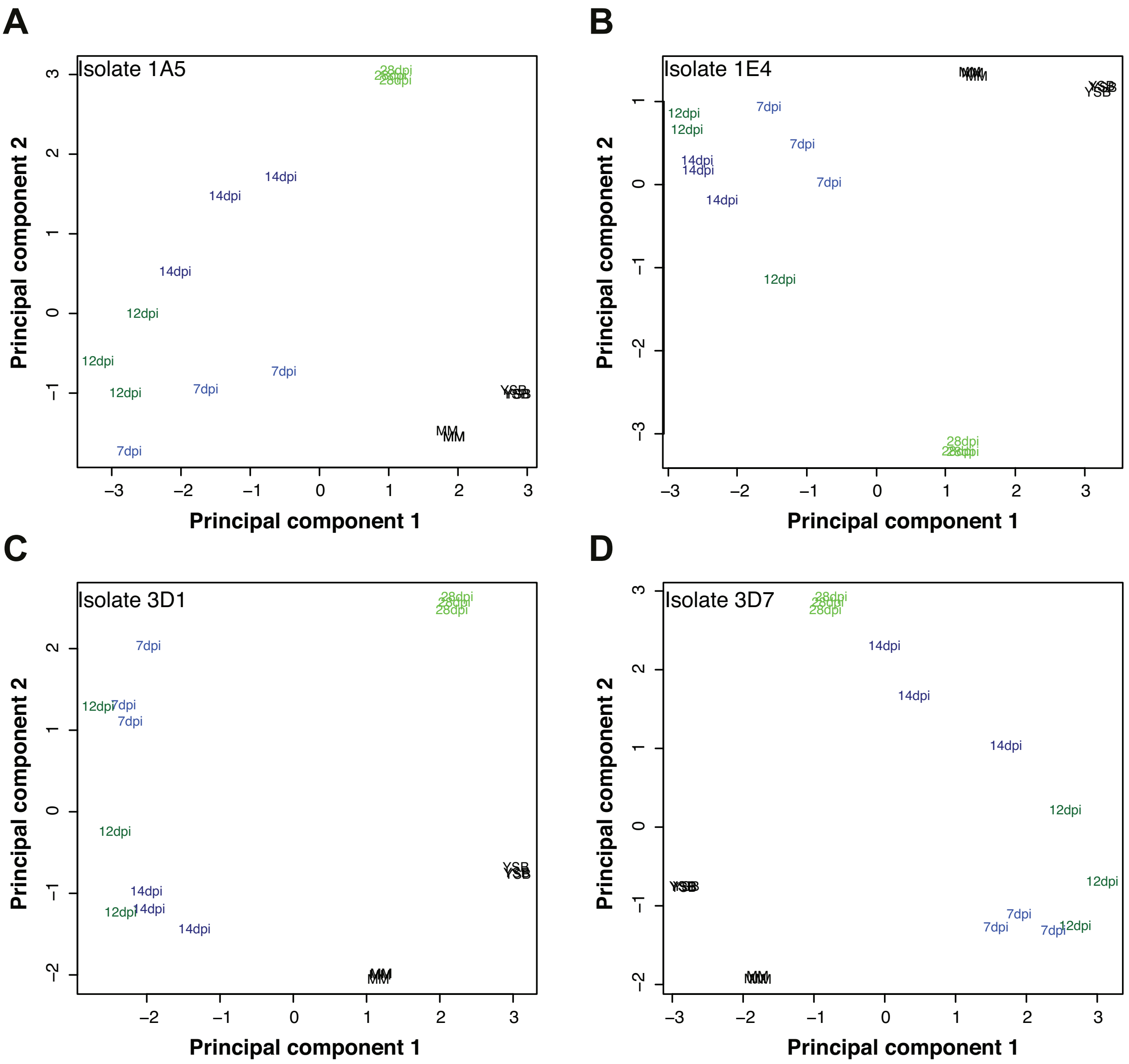


**Supplementary Figure S2: MDS plot of genes and TEs in all four strains.** Biological replicates are shown for each condition and time point.

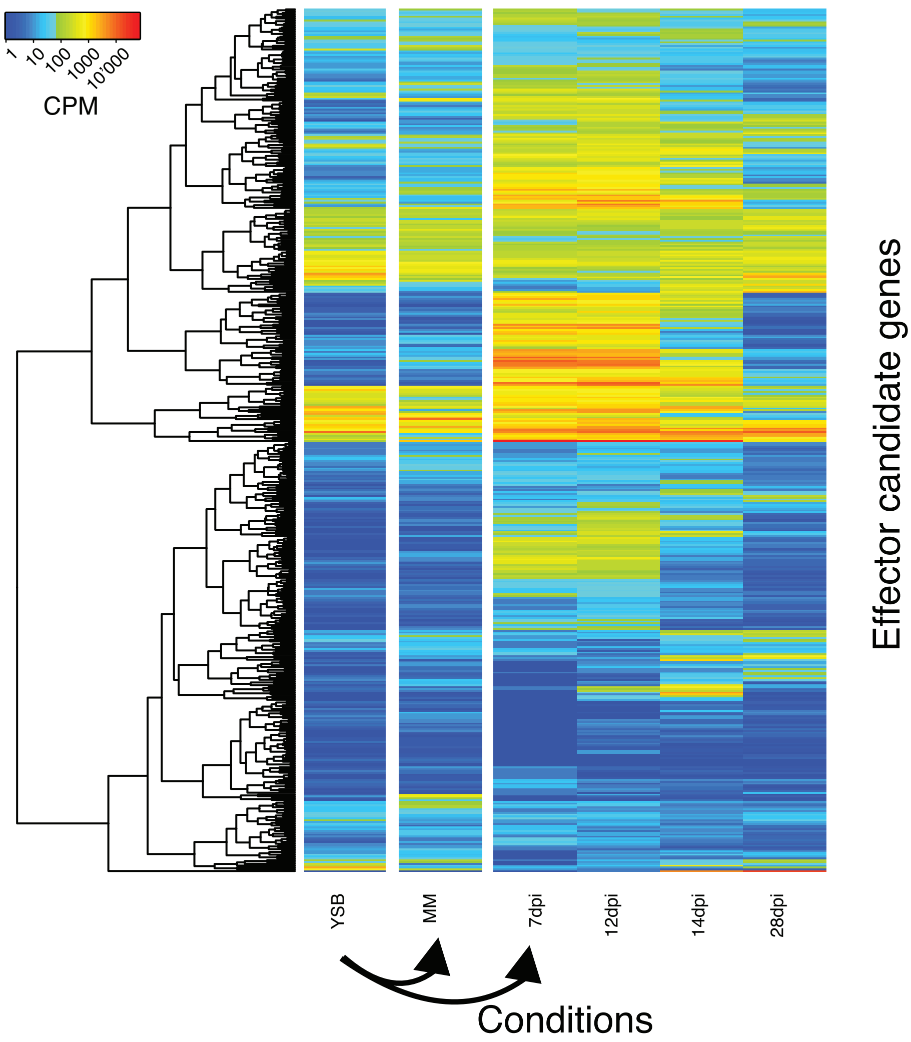


**Supplementary Figure S3: Heatmap of putative effector gene expression in 3D7.**

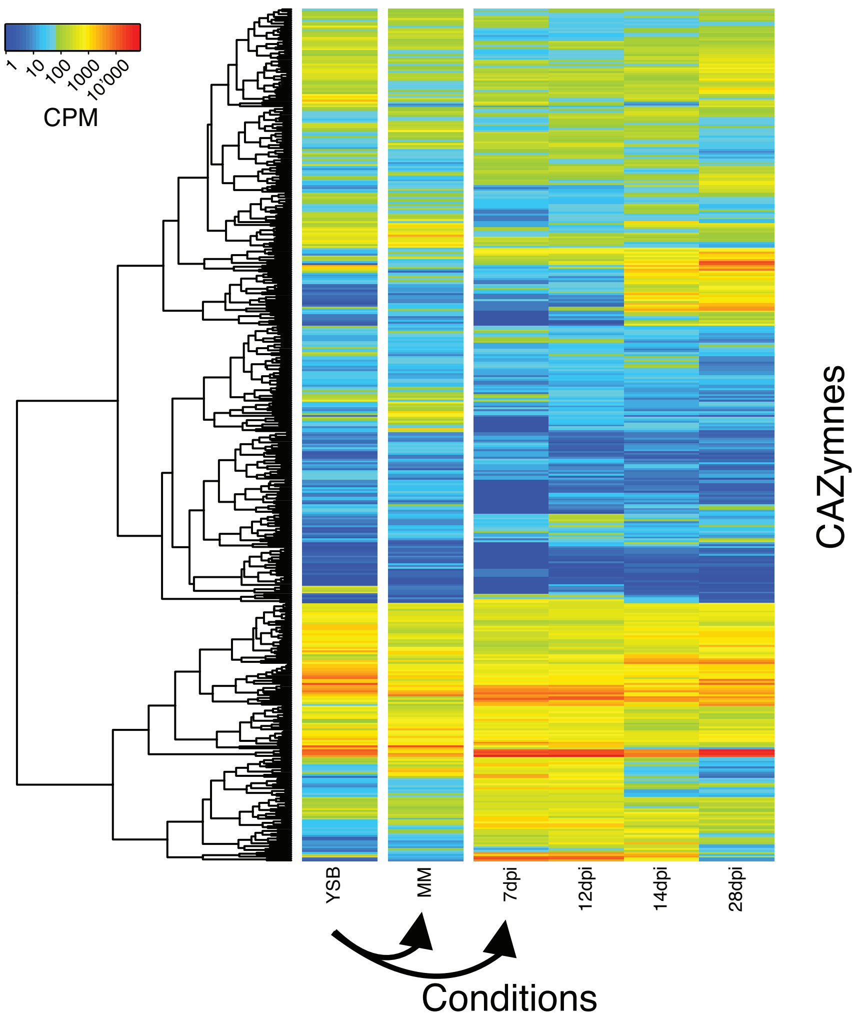


**Supplementary Figure S4: Heatmap of carbohydrate-active enzyme (CAZYme) expression in strain 3D7.**

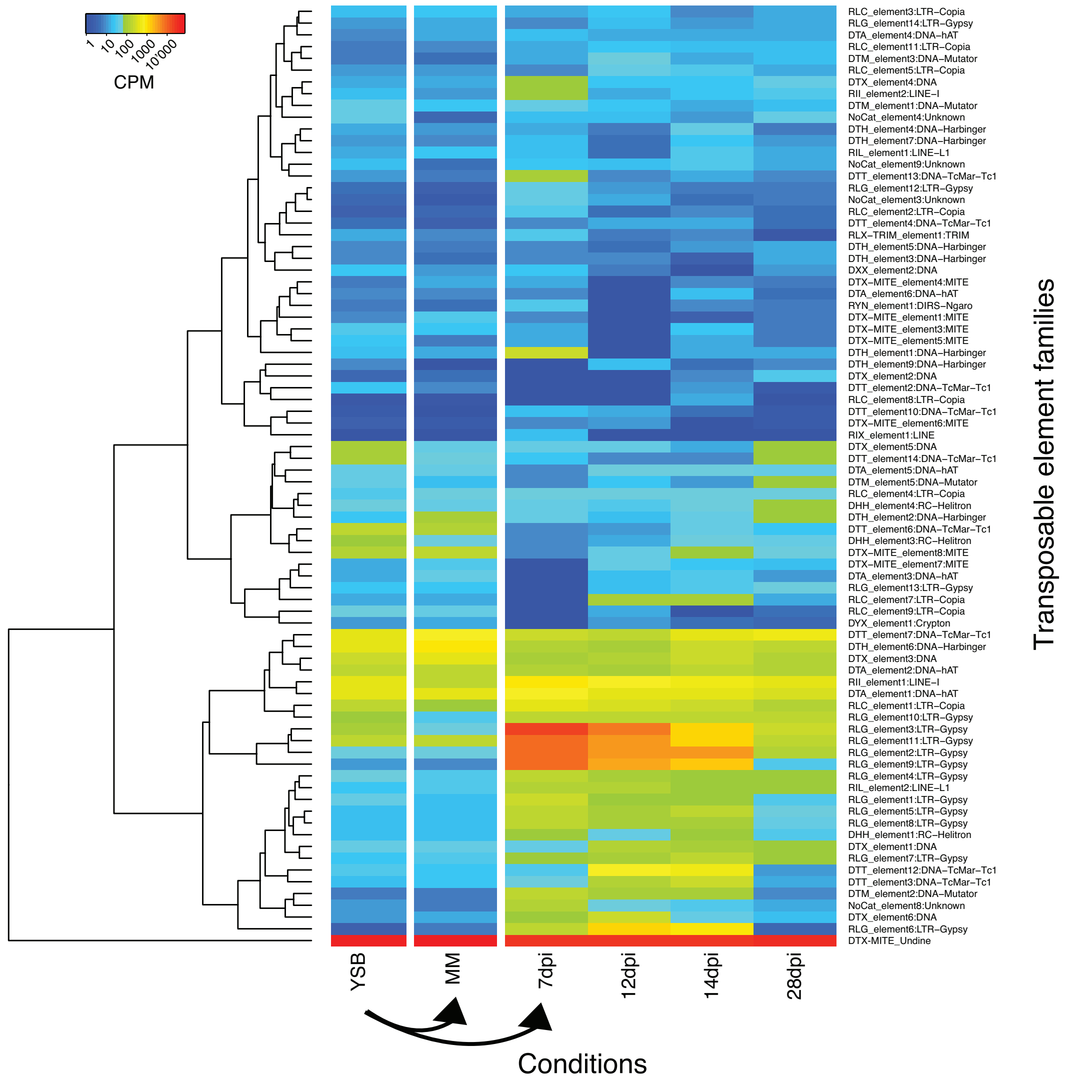


**Supplementary Figure S5A: Heatmap of expression of transposable elements in 3D1**

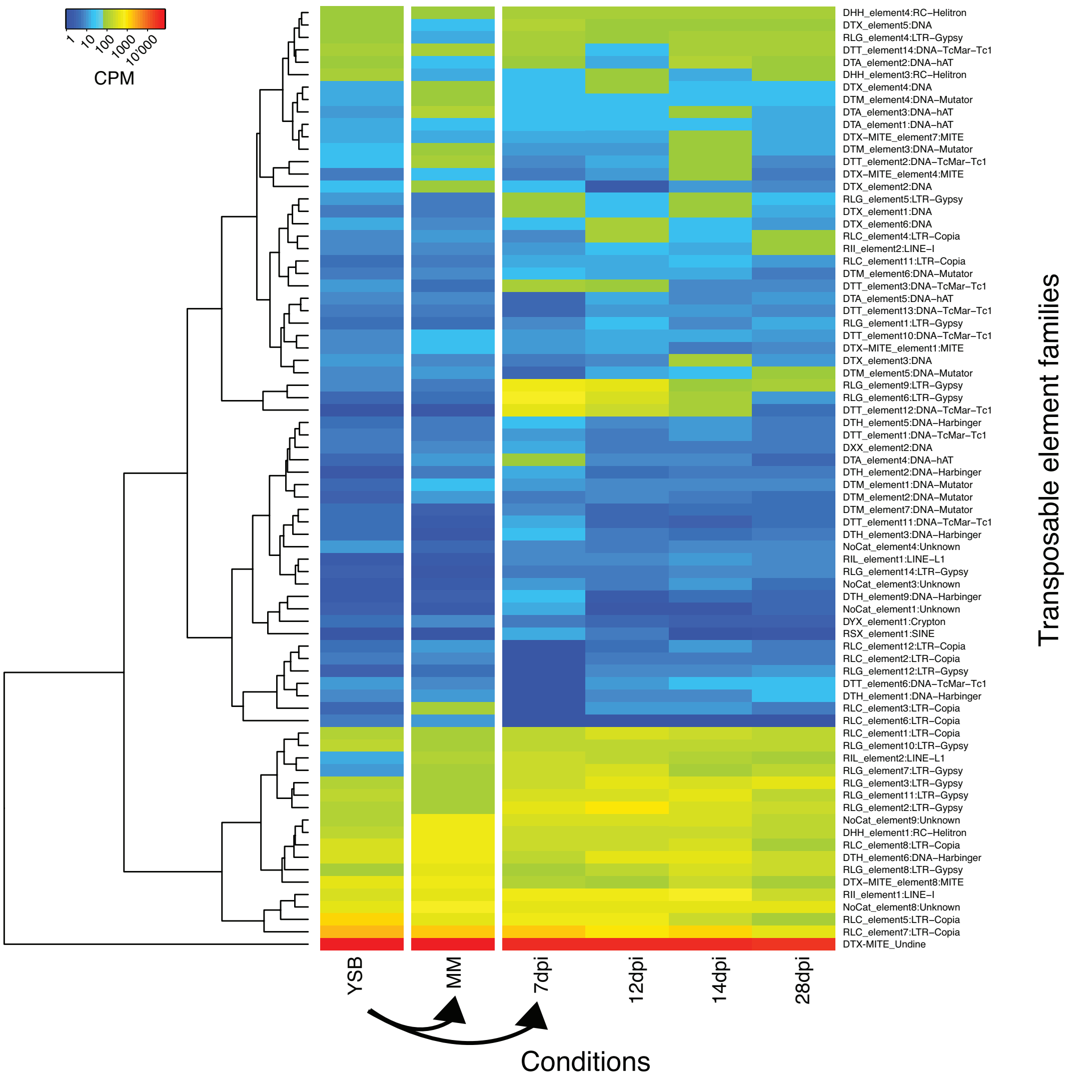


**Supplementary Figure S5B: Heatmap of expression of transposable elements in 3D7**

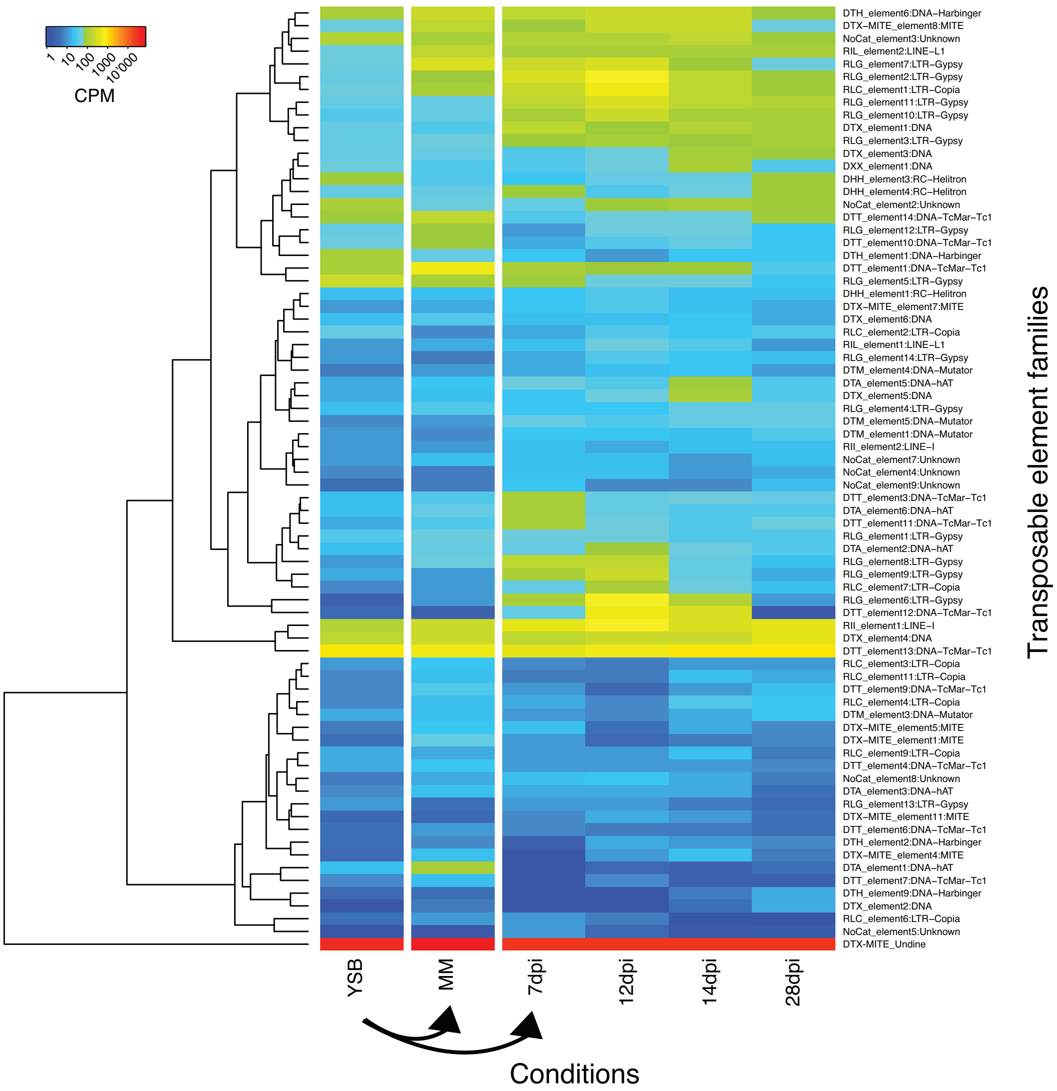


**Supplementary Figure S5C: Heatmap of expression of transposable elements in 1A5**

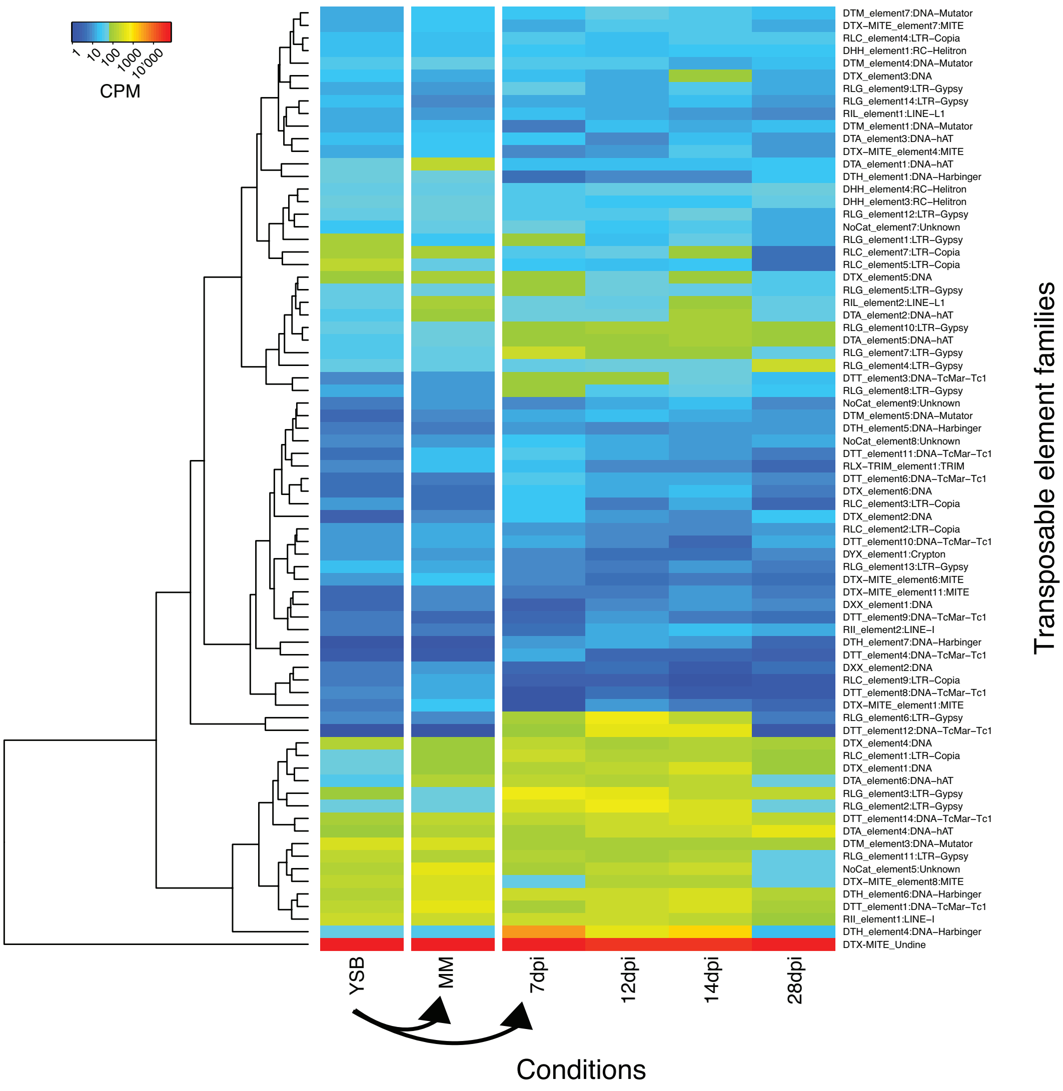


**Supplementary Figure S5D: Heatmap of expression of transposable elements in 3D7**

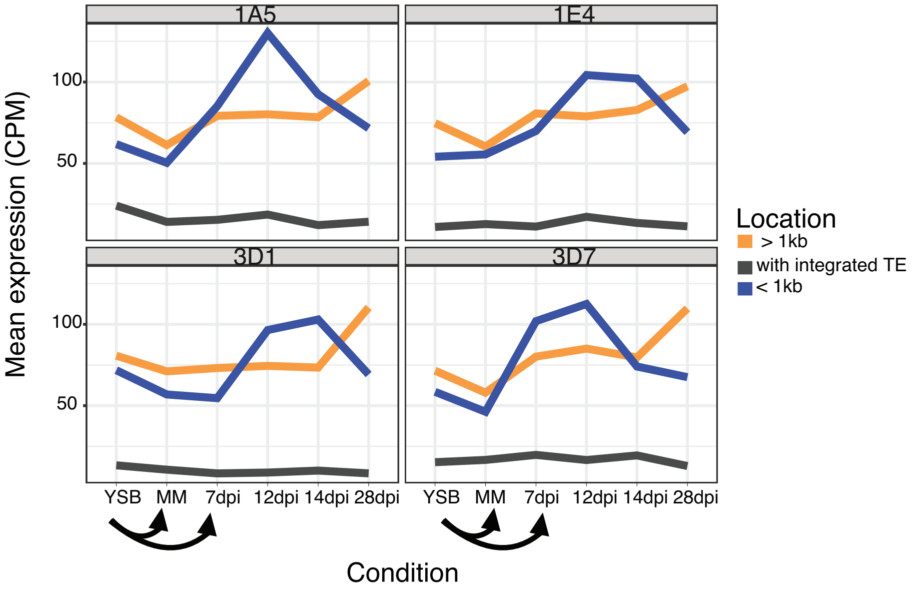


**Supplementary Figure S6: Gene expression as a function of the proximity to transposable elements in all four strains.**

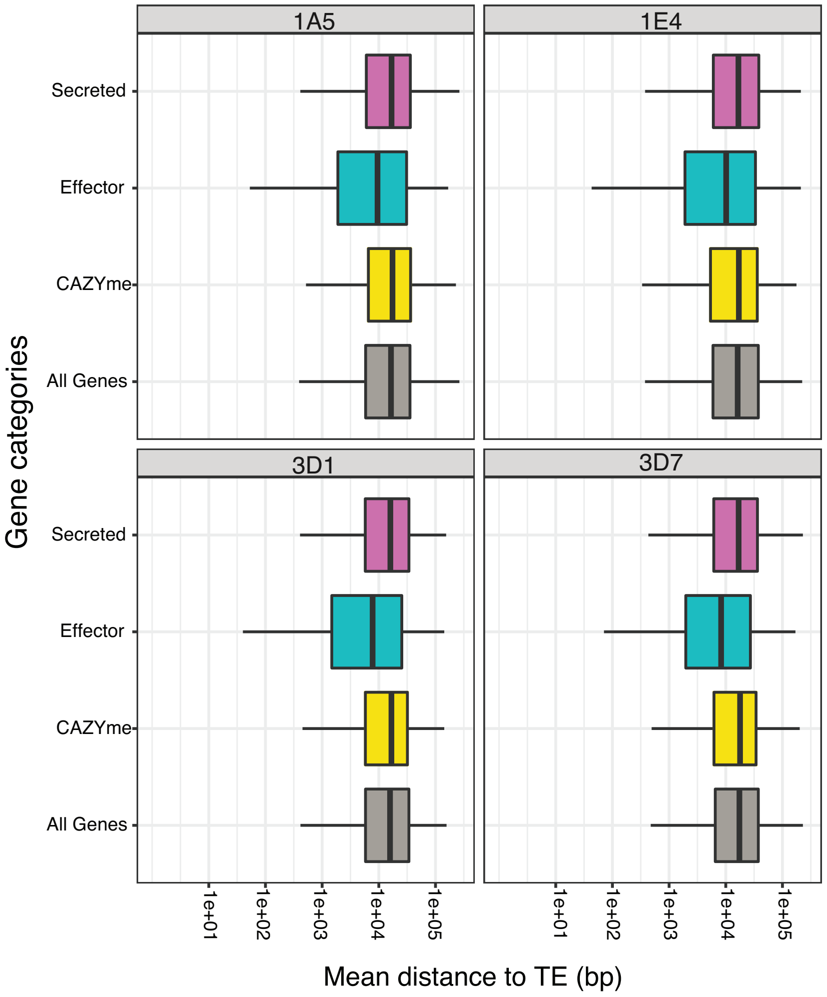


**Supplementary Figure S7: Mean distance between genes grouped by functional category.**

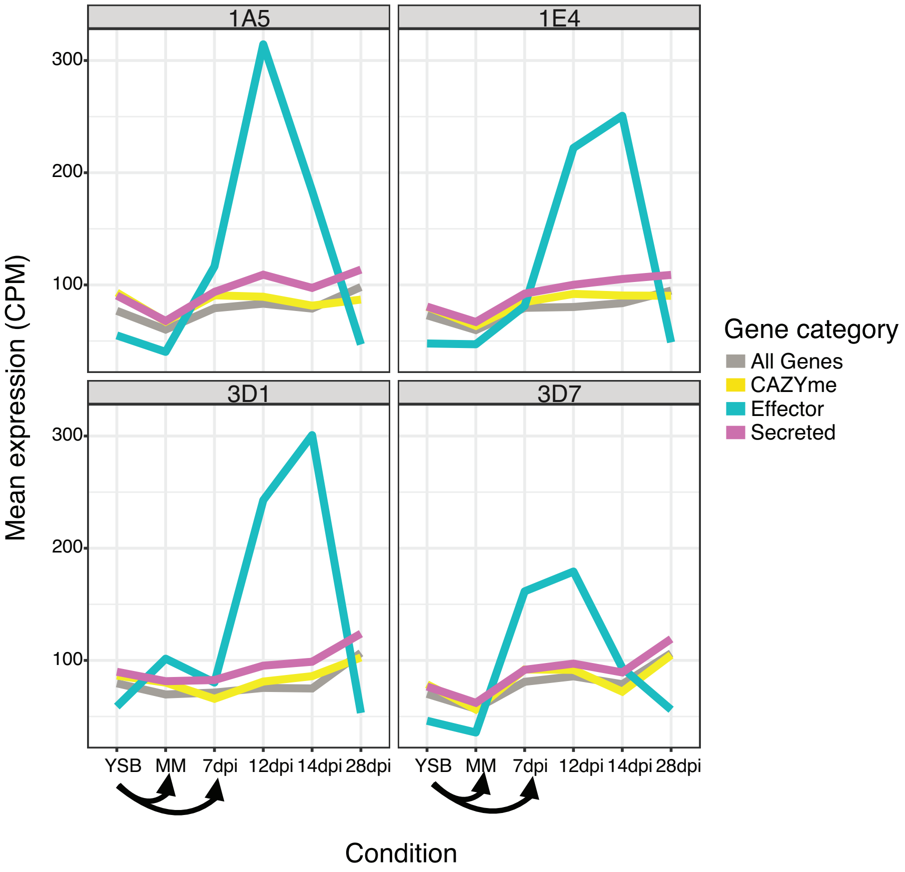


**Supplementary Figure S8: Mean expression of genes grouped by functional category**.

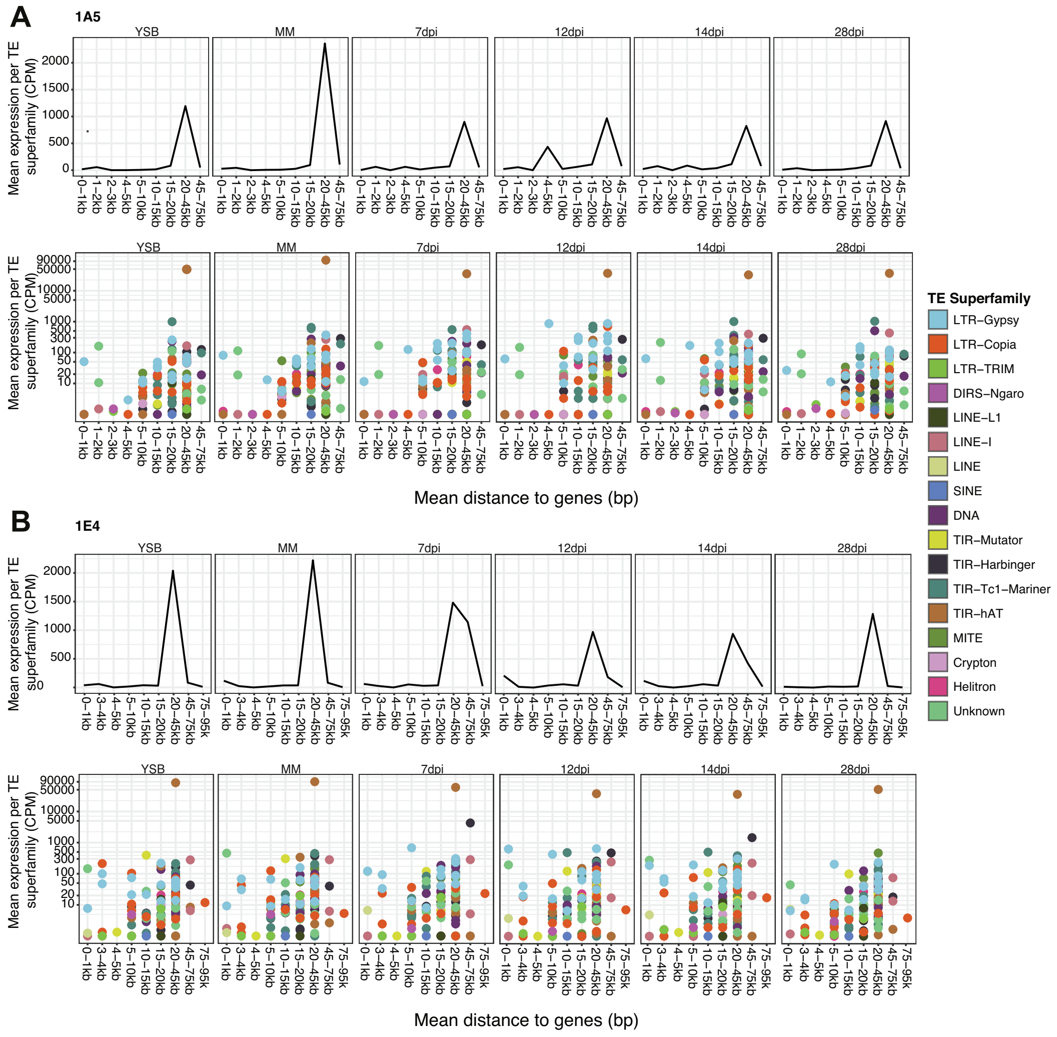


**Supplementary Figure S9: Mean expression of transposable element families at intervals from the nearest gene in 1A5 and 1E4.**

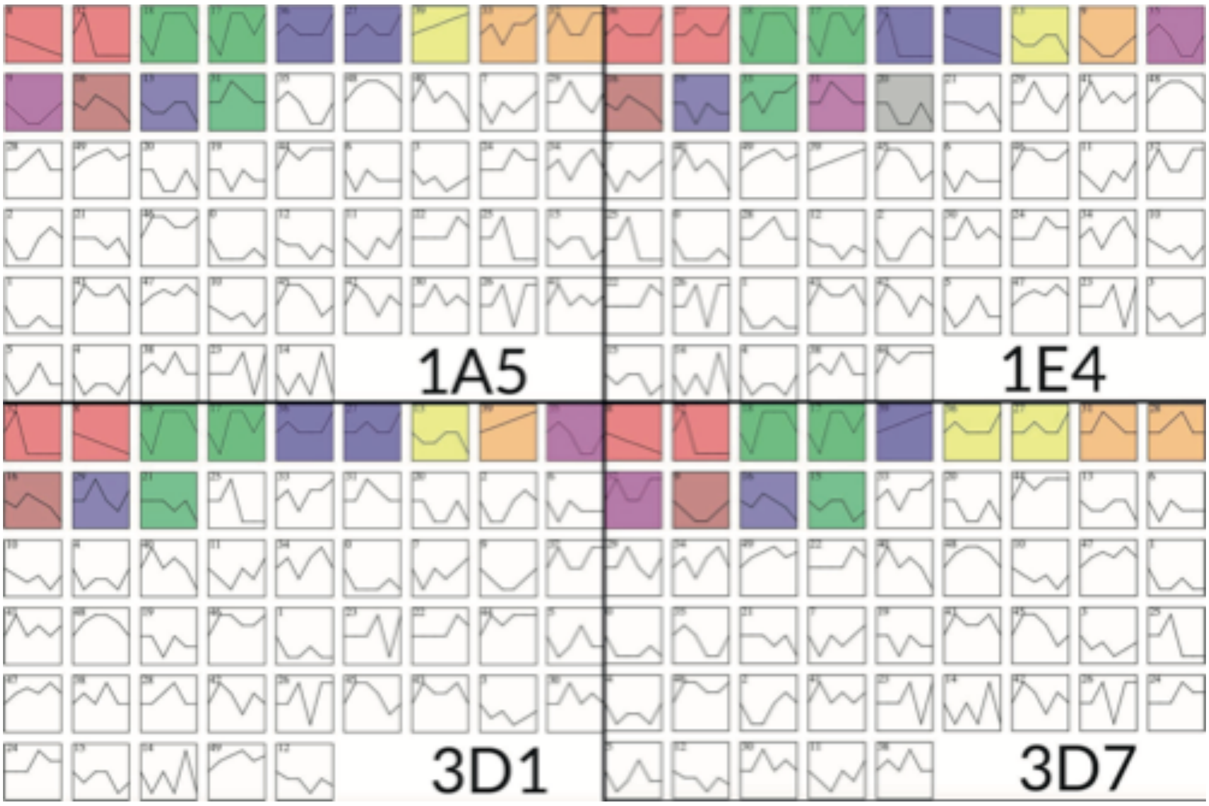


**Supplementary Figure S10: Co-expression profiles for all four strains.** Colored profiles are statistically significant. See methods for details.

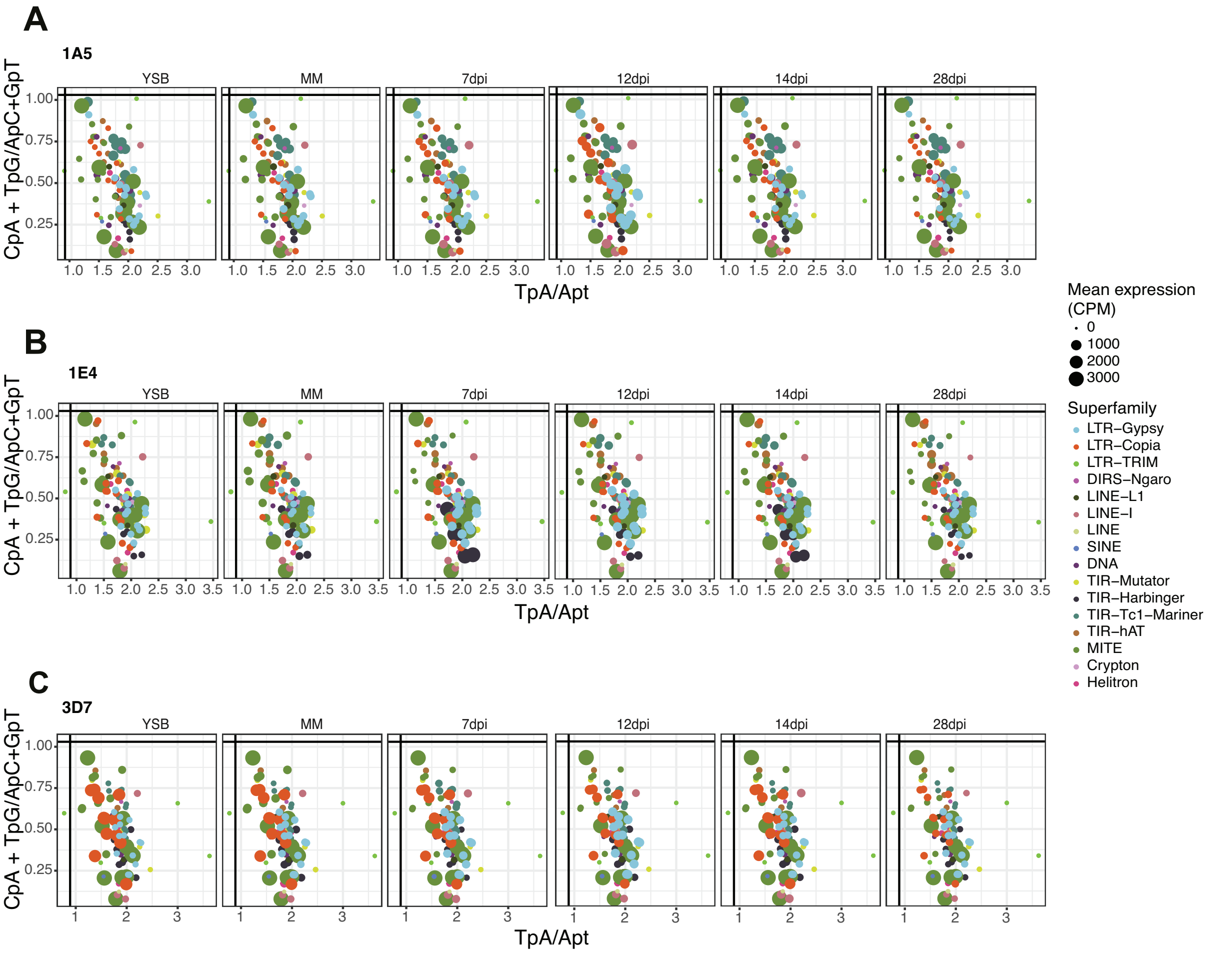


**Supplementary Figure S11:** The RIP indices for each transposable element family and mean expression of the family under all stress conditions for strains 1A5, 1E4 and 3D7. Vertical and horizontal lines represent commonly used thresholds to detect RIP (Hane & Oliver 2008). Colours indicate the superfamily and size the expression at the family level in CPM.

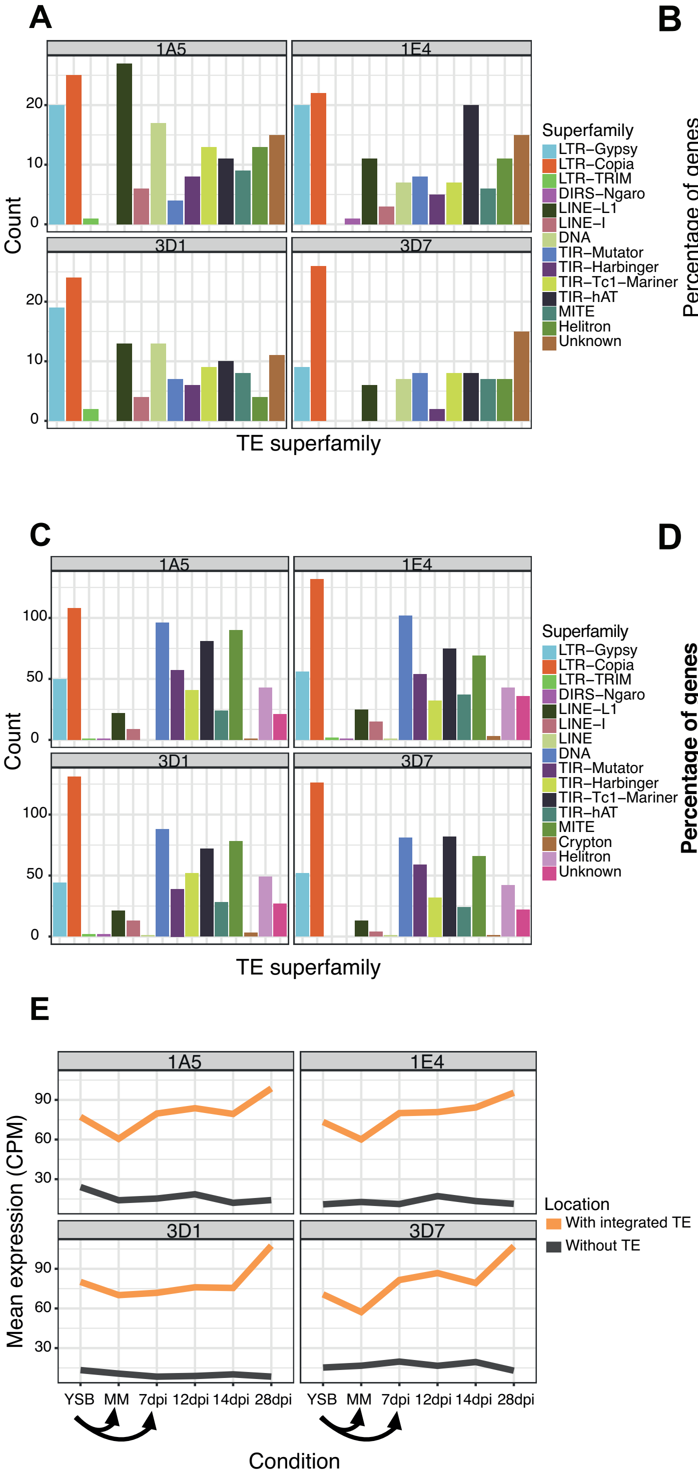


**Supplementary Figure S12:** (A) Transposable element (TE) superfamilies within genes in the four strains. (B) Categories genes with inserted TEs. The percentages are given as the number of genes of that category (core, accessory or strain specific) within genes as a fraction of the total number of genes in a category. (C) TE superfamilies within 1kb of the closest gene. (D) Categories of genes within 1 kb of TEs. The percentages are given as the number of genes of that category (core accessory or strain specific) within genes as a fraction of the total number of genes in a category. (E) Expression of genes with or without inserted TEs.

### Supplementary Tables

(see separate Excel file)

**Supplementary Table S1: Saturation analysis of multiple mapped reads**

**Supplementary Table S2: The average distance of the most highly expressed DTX-MITE-Undine element family from the nearest gene in all four strains**

**Supplementary Table S3: Gene ontology terms of genes in co-expression profiles**

**Supplementary Table S4: The mean distance of transposable element superfamilies to the closest gene**

**Supplementary Table S5: Numbers of transposable element superfamilies within genes**

**Supplementary Table S6: Numbers of transposable element superfamilies within 1 kb of genes**
